# Convergent loss of chemoreceptors across independent origins of slave-making in ants

**DOI:** 10.1101/2021.05.11.443570

**Authors:** Evelien Jongepier, Alice Séguret, Anton Labutin, Barbara Feldmeyer, Claudia Gstöttl, Susanne Foitzik, Jürgen Heinze, Erich Bornberg-Bauer

## Abstract

The evolution of an obligate parasitic lifestyle often leads to the reduction of morphological and physiological traits, which may be accompanied by loss of genes and functions. Slave-maker ants are social parasites that exploit the work force of closely related ant species for social behaviours such as brood care and foraging. Recent divergence between these social parasites and their hosts enables comparative studies of gene family evolution. We sequenced the genomes of eight ant species, representing three independent origins of ant slavery. During the evolution of eusociality, chemoreceptor genes multiplied due to the importance of chemical communication in societies. We investigated evolutionary patterns of chemoreceptors in relation to slave-making in ants. We found that slave-maker ant genomes harboured only half as many gustatory receptors as their hosts, potentially mirroring the outsourcing of foraging tasks to host workers. In addition, parasites had fewer odorant receptors and their loss shows patterns of convergence across origins of parasitism, representing a rare case of convergent molecular evolution. This convergent loss of specific odorant receptors suggests that selective deprivation of receptors is adaptive. The 9-exon odorant receptor subfamily, previously linked to social evolution in insects, was significantly enriched for convergent loss across the three origins of slavery in our study, indicating that the transition to social parasitism in ants is accompanied by the loss of receptors that are likely important for mediating eusocial behaviour. Overall, gene loss in slave-maker ants suggests that a switch to a parasitic lifestyle accompanies relaxed selection on chemical perception.

## INTRODUCTION

Gene losses are pervasive throughout the animal kingdom [Guijarro-Clarke et al., 2020] and may constitute a frequent adaptive evolutionary response [Albalat and Cañestro, 2016]. Indeed, the less-is-more hypothesis [Olson, 1999] proposes that adaptive loss of gene function may occur often and spread through populations, as a change in environment or behaviour can render certain genes non-essential. Additionally, gene loss can open alternative evolutionary trajectories through modifications of genetic network structure, and can even lead to higher fitness in certain environments [Helsen et al., 2020]. Multiple studies have reported cases of convergent gene losses across independent lineages. Examples are the loss of an enzyme required for vitamin C biosynthesis in several vertebrate lineages [Drouin et al., 2011] and the convergent loss of *Paraoxonase 1* in several marine mammals [Meyer et al., 2018]. The latter is likely due to parallel shifts in lipid metabolism in marine ancestors. Such convergent patterns of gene loss accompanying shifts in environment or behaviour suggest that losses may be adaptive. They may reveal which genes are essential in specific environments and which genomic changes underpinned major evolutionary transitions.

One such evolutionary transition and arguably the most common major shift in life history strategy [Poulin and Randhawa, 2015] is the evolution of parasitism. Since parasites exploit their hosts, they often lose the capacity for independent resource acquisition [Sun et al., 2018, Mitreva et al., 2011, Baxter et al., 2010, Kirkness et al., 2010]. Several scenarios may explain the possible benefits of gene losses accompanying the loss of traits during such a transition to parasitism. First, gene losses may release some epistatic constraints because other genes may become free to adapt if those which constrain their activity are lost [Helsen et al., 2020]. Second, losses themselves may change a trait such that a parasite’s fitness is increased, for example because the lost trait itself was detrimental (but not prohibitive) for exploiting the host [Sokurenko et al., 1999]. Finally, losses of genes underlying traits that are dispensable in a new environment (*e.g*. the shift to parasitic lifestyle) may be favored by selection for a reduction in metabolic costs [Albalat and Cañestro, 2016].

While the benefits of acquired genes and the utilisation of existing or duplicated genes [Kondrashov, 2012, Qian and Zhang, 2014] for novel traits have been well studied through comparative evolutionary genomics and transcriptomics [Zhou et al., 2019, Whitelaw et al., 2020, Bernarda, 2020], the possible adaptive benefits of gene losses have only been inferred from parallel morphological trait evolution. Indeed, technical and phylogenetic limitations make it difficult to elucidate the accompanying molecular patterns. Lost genes can no longer be analysed and host-parasite systems which are amenable to experimental or computational investigations typically consist of phylogenetically distant species. To overcome these limitations, we investigated genomic changes underlying the transition to social parasitism in slave-maker ants, an iconic group of social insects already mentioned by Darwin [1859]. Although all ants are ancestrally eusocial, strongly relying on chemical signals to organise tasks within and outside the social colony [Hölldobler, 1995], some species have secondarily lost key social traits, such as the worker caste in inquiline social parasites, or, to a lesser extent, foraging and nursing behaviours in workers of slave-maker ants [Buschinger, 1986].

Slave-maker ants are obligate social parasites that completely rely on workers of closely related host species for brood care, nest defence, foraging and other nest maintenance tasks, all of which demand chemical communication mediated by olfaction. Furthermore, worker reproduction is uncommonly prevalent in slave-maker ants, contrary to their closely related host, suggesting that slave-maker ant workers may have lost their ability to perceive and respond to queen pheromones [Heinze, 1996]. We expect gene losses, specifically the loss of chemosensory genes which radiated during social evolution in ants, to act as an important mechanism underpinning the loss of social behaviour in slave-maker ants.

The myrmicine *Formicoxenus* species group (formerly Formicoxenini, [Blaimer et al., 2018]) is a hot spot for the evolution of social parasitism, with at least five independent origins of slave-making [Beibl et al., 2005, Feldmeyer et al., 2017, Prebus, 2017]. Like most social parasites [Emery, 1909], these slave-maker ants from the genera *Temnothorax* and *Harpagoxenus* exploit closely related non-parasitic species of *Temnothorax* and *Leptothorax* [Hölldobler and Wilson, 1990], although not all members of these taxa are parasitised. The close relatedness between hosts and slave-maker ants and the associated similar genomic architecture render them ideal systems to study co-evolutionary arms races [Foitzik et al., 2001, Feldmeyer et al., 2017, Alleman et al., 2018]. Furthermore, slave-maker ants and hosts share the same nest and thus have very similar ecological and physiological requirements. This reduces possible confounding factors. In addition, closely related non-host, non-parasitic ant species resemble the most likely ancestral state and thus serve as convenient natural controls in genomic comparisons.

In this study, we concentrate on reconstructing the evolutionary history of chemoreceptor (odorant and gustatory) genes and investigate convergent evolutionary patterns across the independent origins of slave-making in ants. We ask: 1) what are the defining changes in chemosensory gene repertoire underlying the repeated evolution of slave-making in ants (specifically, gene losses in these dulotic parasites), 2) are these changes convergent across multiple origins of slave-making in ants, and 3) if so, are these convergent changes (*e.g*. losses) more frequent than expected by chance? The convergent contraction of these putative gene families would indicate parallel evolutionary changes during a shift towards a parasitic lifestyle and underscore a possible adaptive value of these losses. To address these questions, we analyse high quality genome assemblies of eight ant species based on PacBio long read sequencing technology. These include the genomes of three slave-maker ants, representing three independent and distant origins of social parasitism within the *Formicoxenus* group; three hosts, which are the primary hosts of the sequenced slave-maker ant species; and two non-host species as outgroups (see Figure 1).

**Figure 1.**
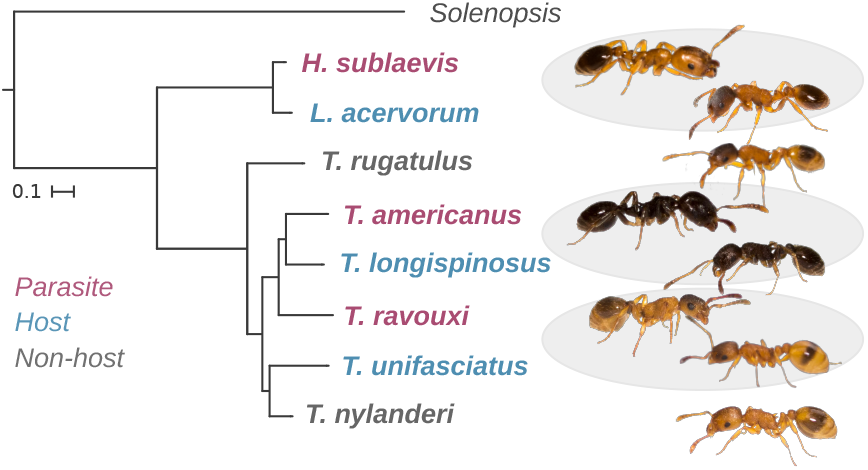
Phylogeny of the focal species in this study. Circles highlight the three independent parasite-host pairs studied. Figure based on Prebus [2017]. Pictures by Barbara Feldmeyer.

## RESULTS

### Eight newly sequenced ant genomes harbour many chemosensory receptor genes

Using a combination of newly sequenced PacBio reads and Illumina short read genomic data for three slave-maker ant species (*Harpagoxenus sublaevis, Temnothorax ravouxi* - formerly *Myrmoxenus ravouxi* -, *Tem-nothorax americanus* - formerly *Protomognathus americanus*), three host species (*Leptothorax acervorum, Temnothorax unifasciatus, Temnothorax longispinosus*) and two non-host outgroup species (*Temnothorax rugatulus, Temnothorax nylanderi*), we assembled eight novel genomes across three independent origins of slave-making (see Methods for details on sample collection, sequencing and genome assembly). Several genome assembly strategies were explored, resulting in eight highly complete (complete `BUSCO`s: mean *±* S.D. = 98.1% *±* 1.2) and comparable genomes (Table 1). We manually annotated 3 718 chemosensory receptor genes across all eight species, including 3 007 odorant receptor genes (*Or* s) and 711 gustatory receptor genes (*Gr*s).

**Table 1:** Genome assembly statistics.

|  | *H. sublaevis* | *L. acervorum* | *T. ravouxi* | *T. unifasciatus* | *T. americanus* | *T. longispinosus* | *T. rugatulus* | *T. nylanderi* |
| --- | --- | --- | --- | --- | --- | --- | --- | --- |
| Estimated genome size | 356 771 525 | 371 301 062 | 369 915 376 | 313 107 076 | 306 525 318 | 309 617 673 | 334 549 751 | 304 816 793 |
| Assembly length | 341 523 168 | 344 708 930 | 362 408 415 | 314 490 334 | 279 543 510 | 298 344 416 | 355 735 536 | 313 389 767 |
| GC (%) | 39.00 | 38.67 | 38.70 | 38.81 | 39.20 | 38.93 | 39.31 | 39.06 |
| Number of contigs | 2 750 | 1 269 | 4 380 | 2 496 | 1 229 | 3 430 | 2 079 | 1 555 |
| Largest contig | 4 363 893 | 6 338 531 | 1 282 434 | 2 901 263 | 4 381 237 | 1 041 114 | 4 513 070 | 4 914 139 |
| N50 | 391 101 | 730 108 | 128 698 | 359 731 | 603 026 | 156 499 | 450 769 | 619 324 |
| L50 | 176 | 104 | 751 | 196 | 121 | 541 | 155 | 111 |
| Complete BUSCOs (%) | 98.1 | 98.3 | 95.7 | 98.2 | 98.2 | 95.6 | 98.4 | 98.5 |
| Duplicated BUSCOs (%) | 1.4 | 1.6 | 7.7 | 2.8 | 1.3 | 1.8 | 2.3 | 2.4 |
| Illumina mapping rate (%) | 97.87 | 97.24 | 96.76 | 96.32 | 97.22 | 95.77 | 95.98 | 97.33 |
| PacBio mapping rate (%) | 91.89 | 91.59 | 96.28 | 91.24 | 97.34 | 92.11 | 94.08 | 93.27 |

### Extensive chemosensory receptor losses in slave-maker ants, modest expansions in hosts

In order to identify *Or* and *Gr* losses or expansions in the eight genomes, we first identified orthologous clusters of *Or* s and *Gr*s by taking an explicit phylogenetic approach (see Methods, Fig. 2 A1 and Fig. 3 A). Our approach to identify orthologs using not only three host/slave-maker replicates but also multiple outgroup species to differentiate alternative evolutionary scenarios, combined with the manual verification of each putative chemoreceptor annotation, represents the most detailed chemoreceptor analysis to date.

**Figure 2.**
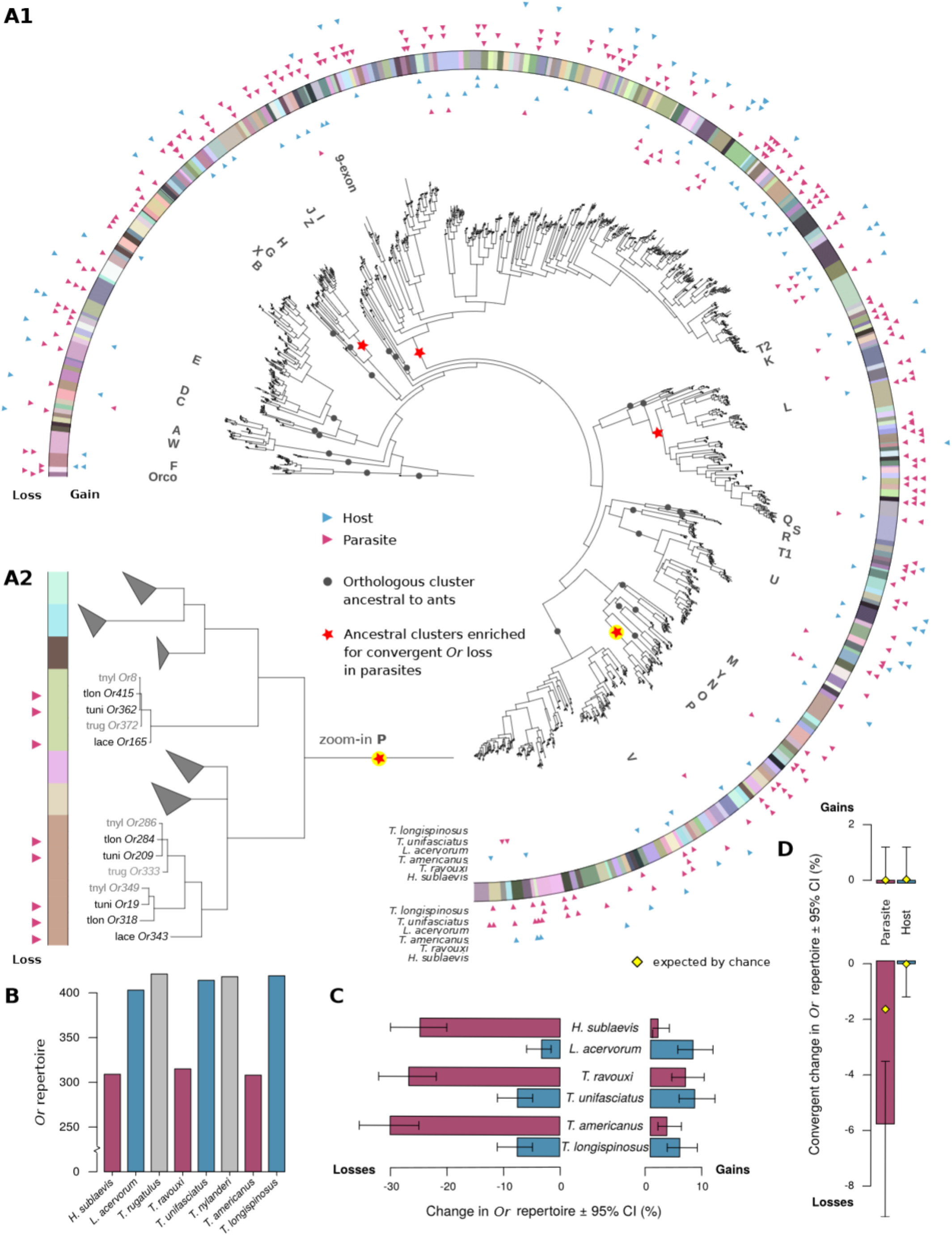
Distribution of odorant receptor gene (*Or*) gains and losses across species and orthologous clusters. A1: Phylogenetic clustering of *Or* s across three slave-maker ant species and their hosts. Species-specific expansions were collapsed for visualisation purposes. Orthologous cluster assignments are displayed through colours in the outer ring. Subfamilies identified in previous studies as being ancestral to the ants, bees and wasps are represented by dark circles and corresponding letters throughout the tree. We found that four of these ancestral clusters were enriched for convergent *Or* loss in parasites, marked by the red stars in the tree. Triangles on the inner side of the coloured ring show the distribution of gene gains across orthologous clusters and triangles on the outer side of the ring show the distribution of gene losses. Species names at the end of the ring indicate the species in which the gains or losses occurred, with red triangles showing gains/losses in slave-maker ant (parasite) species and blue triangles showing gains/losses in host species. A2: Zoom-in on ancestral cluster P, containing seven orthologous clusters. Five clusters, in which no genes were gained or lost, were collapsed. In the remaining two clusters, gene copies were convergently lost in all three slave-maker ant species. Red triangles indicate a gene loss, as exhibited by the presence of a copy in a host species, but not in its respective parasite species. B: Number of *Or* copies in each focal species. Bars are colour-coded according to lifestyle of the species as in the following graphs, with red representing parasite species, blue host species and grey outgroup species. C: Relative change in *Or* repertoire in slave-maker ants compared to their respective hosts, and in hosts compared to their respective parasites. D: Number of cases of convergent gain (top) or loss (bottom) of *Or*s in slave-maker ants and in hosts. An event (gain or loss) was defined as convergent if it occurred in all three parasite or host species within an orthologous cluster. The yellow diamonds represent the null-probability, set to the product of the marginal probabilities of loss for each parasite, *i.e*. the probability that convergent loss occurred even though losses are completely independent in each of the three parasites.

**Figure 3.**
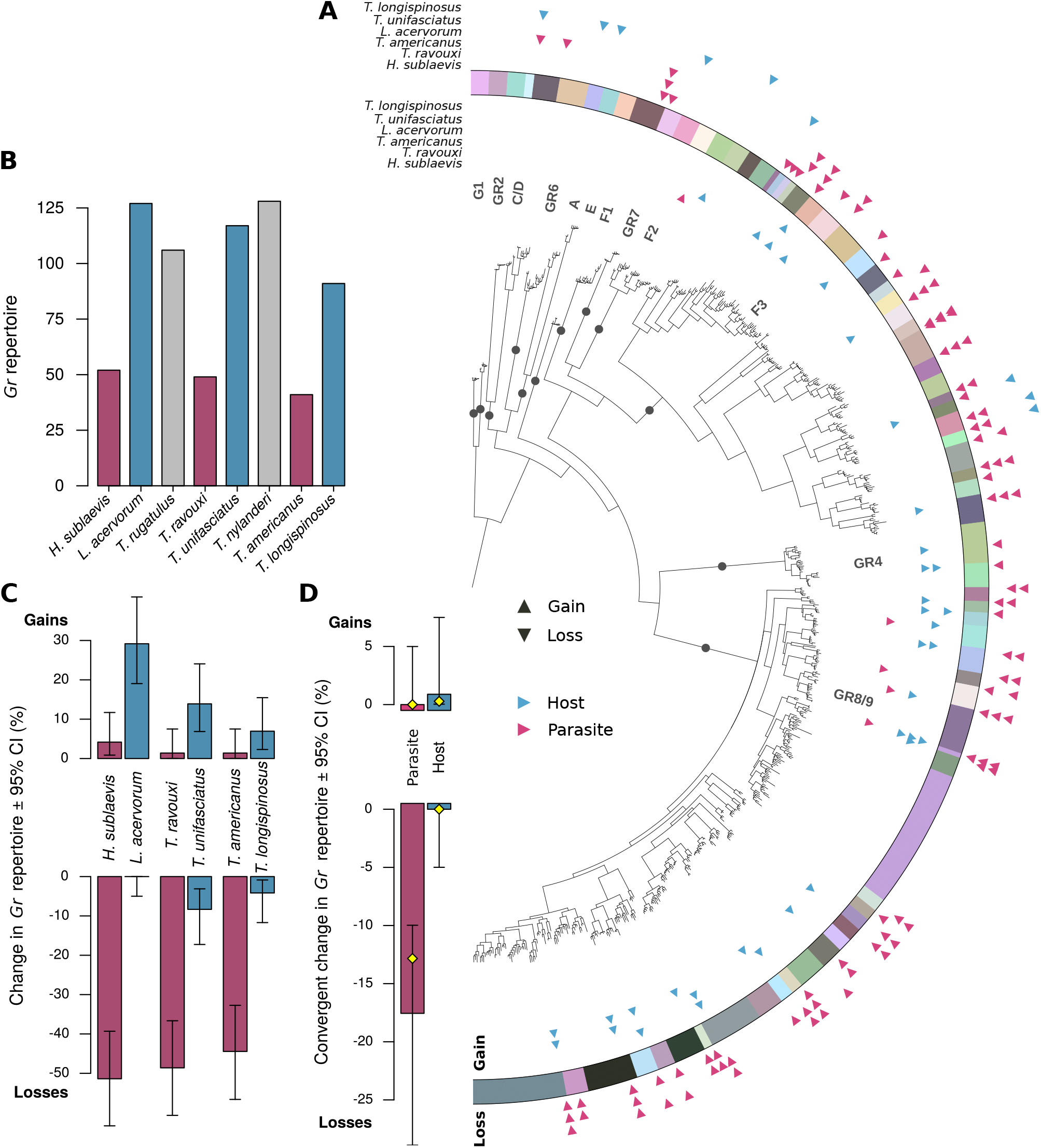
Distribution of gustatory receptor gene (*Gr*) gains and losses across species and orthologous clusters. A: Phylogenetic clustering of *Gr*s across three slave-maker ant species and their hosts. The gene tree was constructed using the same methods described in Figure 2. Orthologous cluster assignment is displayed through colours in the outer ring. Subfamilies identified in previous studies as being ancestral to the ants, bees and wasps are represented by dark circles and corresponding letters throughout the tree. Triangles on the inner side of the coloured ring show the distribution of gene gains across orthologous clusters and triangles on the outer side of the ring show the distribution of gene losses. Species names at the top indicate the species in which the gains or losses occurred, with red triangles showing gains/losses in slave-maker ant (parasite) species and blue triangles showing gains/losses in host species. B. Number of *Gr* copies in each focal species. Bars are colour-coded according to lifestyle of the species as in following graphs, with red representing parasite species, blue host species and grey outgroup species. C. Relative change in *Gr* repertoire in slave-maker ants compared to their respective hosts, and in hosts compared to their respective parasites. D: Number of cases of convergent gain (top) or loss (bottom) of *Gr*s in slave-maker ants and in hosts. An event (gain or loss) was defined as convergent if it occurred in all three parasite or host species within an orthologous cluster. The yellow diamonds represent the null-probability, set to the product of the marginal probabilities of loss for each parasite, *i.e*. the probability that convergent loss occurred even though losses are completely independent in each of the three parasites.

To determine where a gene loss had occurred in a slave-maker ant species, for each orthologous cluster we assessed whether slave-maker ant species had fewer chemosensory receptor genes than their hosts and outgroup species. For one-to-one orthologs, this means that a host’s single copy was lost in its parasite. For partial out-paralogs (*i.e*. those assigned to the same orthologous cluster because the duplication event occurred in the common ancestor of some, but not all of our focal species), this translates into a parasite having lost one or more out-paralogs compared to its host and outgroup species (Supplementary Figure S3). To determine where a gene gain had occurred, we identified orthologous clusters which only contained a single gene copy from each of the host and outgroup genomes, but which contained multiple gene copies in one or several parasite genomes. This indicates the gain of one or several in-paralogs in such parasite species since divergence from their host. The same analyses were conducted for the hosts in comparison to the slave-maker species.

Slave-maker ants exhibited smaller *Or* repertoires compared to their respective host species and both non-host species (Fig. 2 B). Slave-maker ant genomes contained on average 311 *Or* s (range 308-315), and the number of *Or* s did not vary significantly between slave-maker ant species (3-sample test for equality of proportions with continuity correction: *χ*^2^ = 1.838, *d.f*. = 2, *p*_*adj*_ = 0.3990). In contrast, each host and non-host genome harboured more than 400 *Or* s (range 403-421, Supp. Table S11), again with *Or* numbers similar across all host species (3-sample test for equality of proportions with continuity correction: *χ*^2^ = 6.427, *d.f*. = 2, *p*_*adj*_ = 0.080). Accordingly, we found that slave-maker ants experienced significantly more *Or* losses than their respective hosts across all three origins of parasitism (Fig. 2 C; Supplementary Table S12; 2-sample test for equality of proportions with continuity correction, *L. acervorum*/*H. sublaevis*: *χ*^2^ = 58.34, *d.f*. = 1, *p*_*adj*_ = 6.6330 *×* 10^*-*14^; *T. unifasciatus*/*T. ravouxi* : *χ*^2^ = 41.81, *d.f*. = 1, *p*_*adj*_ = 1.007 *×* 10^*-*10^; *T. longispinosus*/*T. americanus*: *χ*^2^ = 49.48, *d.f*. = 1, *p*_*adj*_ = 3.014 *×* 10^*-*12^). Slave-maker ants also exhibited smaller *Gr* repertoires than their respective hosts across all three origins of parasitism (parasite range 41-52; host and non-host range 91-128; Fig. 3 B), with *Gr* numbers similar across parasite genomes on one hand (3-sample test for equality of proportions with continuity correction: *χ*^2^ = 1.927, *d.f*. = 2, *p*_*adj*_ = 0.3816) and across host genomes on the other hand (3-sample test for equality of proportions with continuity correction: *χ*^2^ = 6.261, *d.f*. = 2, *p*_*adj*_ = 0.087). Accordingly, significantly more *Gr* losses were found in the genomes of slave-maker ants than in the genomes of their respective hosts (Fig. 3 C; Supplementary Table S12; 2-sample test for equality of proportions with continuity correction, *L. acervorum*/*H. sublaevis*: *χ*^2^ = 52.65, *d.f*. = 1, *p*_*adj*_ = 1.1958 *×* 10^*-*12^; *T. unifasciatus*/*T. ravouxi* : *χ*^2^ = 25.24, *d.f*. = 1, *p*_*adj*_ = 5.0760 *×* 10^*-*7^; *T. longispinosus*/*T. americanus*: *χ*^2^ = 29.59, *d.f*. = 1, *p*_*adj*_ = 7.9965 *×* 10^*-*8^). This indicates that the transition to parasitism is associated with extensive losses of both *Or* s and *Gr*s in the parasites’ genome.

Host species, on the other hand, exhibited modest species-specific expansions of their *Or* repertoires, but the magnitude of these expansions did not differ from those in slave-maker ant species, with the exception of the species pair *H. sublaevis* and *L. acervorum* (for the proportion of *Or* clusters which underwent expansions in each species, see Supplementary Table S12). *L. acervorum* exhibited significantly more *Or* expansions than its parasite *H. sublaevis* (23 versus 4 clusters; 2-sample test for equality of proportions with continuity correction: *χ*^2^ = 12.55, *d.f*. = 1, *p*_*adj*_ = 0.0012). *Gr* expansions were also moderate across host species, but hosts exhibited significantly more species-specific duplications of *Gr*s compared to their respective parasite for two of the three species pairs in our study: *H. sublaevis* and *L. acervorum* (2-sample test for equality of proportions with continuity correction: *χ*^2^ = 14.45, *d.f*. = 1, *p*_*adj*_ = 4.317 *×* 10^*-*4^), and *T. ravouxi* and *T. unifasciatus* (2-sample test for equality of proportions with continuity correction: *χ*^2^ = 6.299, *d.f*. = 1, *p*_*adj*_ = 0.0181).

### Convergent levels of chemoreceptor loss in slave-maker ants across three origins of parasitism

Slave-maker ants exhibited comparable levels of *Or* loss across all three origins of parasitism, ranging from 25.08% in *H. sublaevis* to 27.69% in *T. ravouxi* and 29.97% in *T. americanus* (Fig. 2 C; Supplementary Table S12). Host species on the other hand retained similar numbers of *Or* s compared to the non-host species. Accordingly, hosts exhibited very low levels of *Or* loss, ranging from 3.26% in *L. acervorum* to 7.49% in *T. unifasciatus* as well as in *T. longispinosus*.

Similarly, the proportion of *Gr*s that were lost in slave-maker ant species compared to their hosts was comparable across the three origins of parasitism, ranging from 55.56% in *H. sublaevis* to 47.22% in *T. ravouxi* and 44.44% in *T. americanus* (Fig. 3 C; Supplementary Table S12). The proportion of *Gr*s lost in host genomes was much lower than in slave-maker ants, ranging from none in *L. acervorum* to 8.33% in *T. unifasciatus* and 4.17% in *T. longispinosus*. Like in parasites, losses did not differ between host species.

### Convergent loss of specific receptors across three origins or parasitism

For both chemoreceptor families, we tested whether convergent loss occurred more often than expected by chance. Convergent loss was defined as the occurrence of chemoreceptor loss in all three slave-maker ant species within an orthologous cluster. In total, 20 out of 307 orthologous odorant receptor clusters showed convergent loss in the three slave-maker ant species (Fig. 2 D), which was more than expected by chance (exact binomial test: expected prob. = 0.0174; observed prob. [95% CI] = 0.0652 [0.0402-0.0988]; *n* = 307; *p* = 6.762 *×* 10^*-*7^; power = 0.9235). In contrast, not a single orthologous cluster underwent convergent loss in the hosts. This is not surprising given that loss was so rare in hosts that the probability that all three hosts lost the same *Or* by chance was only 7.949 *×* 10^*-*5^.

Previous studies on social and solitary Hymenoptera identified 30 *Or* subfamilies, represented by the dark circles and corresponding letters in Figure 2 A1, each one representing one or a few genes that are ancestral to the ants, bees, and wasps [Zhou et al., 2015, 2012]. We find that four of these ancestral clusters, represented by the red stars in Figure 2 A1, were enriched for convergent *Or* loss in slave-maker ants (Table 2). These included the 9-exon subtree, which is the largest subtree in ants and is often implicated in the evolution of eusociality. McKenzie et al. (2016) showed exceptionally high gene birth rates for 9-exon *Or* s, in particular in the ancestors of ants, followed by continued expansions in separate ant lineages. However, none of the ant species studied to date showed particularly high gene death rates. Like the 9-exon *Or* s, the other three subfamilies enriched for convergent *Or* loss have also been subject to gene family expansions in the ants (Zhou et al. 2015, Engsontia et al. 2015), including subtrees H, L and P (Fig. 2).

**Table 2:** *Or* subtree convergent loss statistics (exact binomial test performed for each subtree).

| Subtree | Observed probability | 95% CI | $n$ | $p_{adj}$ |
| --- | --- | --- | --- | --- |
| 9 exon | 0.080 | 0.037-0.147 | 112 | $9.697e - 4$ |
| H | 0.250 | 0.032-0.651 | 8 | 0.0119 |
| L | 0.105 | 0.029-0.248 | 38 | 0.0119 |
| P | 0.286 | 0.037-0.710 | 7 | 0.0119 |

Thirteen out of 72 orthologous gustatory receptor clusters showed convergent loss in the three slave-maker ant species (Fig. 3 D), which was expected by chance (exact binomial test: expected prob. = 0.1457; observed prob. [95% CI] = 0.1806 [0.0998-0.2889]; *n* = 72; *p* = 0.4027). No orthologous cluster underwent convergent loss in the hosts. To allow for comparisons between the two chemoreceptor families despite large differences in the total number of receptors, a power analysis was performed. Assuming a sample size of 307 orthologous clusters for *Gr*s, as is the case for *Or* s in our study, the probability of detecting convergent losses of specific *Gr*s above what is expected by chance was only 35% (*i.e*. power = 0.35 assuming n = 307 orthologous *Gr* clusters). In contrast, the probability of detecting significant convergent losses of specific *Or* s was 92%. This result highlights that the absence of detection of convergent loss for *Gr*s, but not for *Or* s, is not simply a statistical artefact of the lower sample size for *Gr*s. Rather, it reflects a fundamental difference in effect size (level of convergence) for *Gr*s and *Or* s.

## DISCUSSION

The results presented here demonstrate how, even between closely related species, gene losses accompany the evolutionary transition to becoming a parasite of the sister species. While most studies on gene losses in parasites have so far concentrated on the relationship between species which are essentially unrelated (*e.g*. bacteria and their animal [Sokurenko et al., 1999] or plant hosts [Hulin et al., 2018], parasitic plants and their distantly related plant hosts [Sun et al., 2018]), the system studied here evolved over a time span of only 8-28 million years [Prebus, 2017].

We concentrated on chemoreceptors because they are pivotal to ant sociality, the key trait which had facilitated the world-wide success of ants in first place *ca* 100 million years ago [Ward, 2014], and because we expected chemoreceptors to have undergone losses in slave-maker ants alongside the loss of certain social behaviours. Overall, we find smaller repertoires of both *Or* s and *Gr*s in all three slave-maker ant species compared to their respective host species and the outgroups but we found different patterns for *Or* s and *Gr*s with respect to the convergence of these losses.

*Or* s are a hallmark of social insects as they underwent a remarkable gene family expansion in social compared to solitary species. For example, while 66 *Or* s are known in *Drosophila* and 20-92 *Or* s in Lepidopteran genomes, ant genomes feature several hundred *Or* s [Engsontia et al., 2015, Slone et al., 2017]. With around 400 *Or* gene copies in each of the host and non-host species, the genomes presented here harbour among the most elaborate repertoire found in ants (see also McKenzie and Kronauer [2018]). Most remarkably, considering that these ant species build relatively small societies (less than 100 members on average) and lack morphological variation in the worker caste, we find that their *Or* repertoire exceeds even that of the highly social leaf-cutter ants [Engsontia et al., 2015]. However, these high numbers likely reflect the high quality of our PacBio genomes, rather than biological differences and underscore the reliability of our data (see also tables S3-S10). Most importantly, this high quality allows the notoriously difficult inference of absent genomic states (genes).

Furthermore, we find extensive *Or* losses in all three slave-maker ant species, likely reflecting a diminished need to communicate during foraging and brood care tasks. Such a convergent pattern in the extent of *Or* losses suggests that conserved genes, which played an important role in the ancestor of the slave-maker ant species studied here, suddenly lost their function or even became maladaptive.

This is further underscored by the observation of repeated loss of exactly the same ancestral *Or* s in all three parasite lineages (Figure 2 D). This convergence with reference to individual genes suggests that slave-maker ants retain specific receptors that are necessary for the successful identification of host colonies as well as for chemical communication during raids [Regnier and Wilson, 1971], which may translate into convergent loss of the receptors that are not needed for such tasks. Alternatively, this convergence may indicate that the loss of important social traits following the transition to a parasitic life style goes hand in hand with the loss of *Or* s implicated in *specific* traits that are instrumental in their sociality. The 9-exon *Or* subfamily, which has been linked to social signalling and even social evolution in insects [Pask et al., 2017], was significantly enriched for convergent loss in the parasites in our study. This suggests that the transition to slave-making in ants is accompanied by a loss of receptors that are important for mediating eusocial behaviour. Similarly, though to a lesser extent, the convergent loss of *Or* s from the H and P subtrees in slave-maker ants suggests a loss of receptors linked to colony tasks. Indeed, many *Or* s in subtrees H and P show worker-biased expression compared to males in various ant species, as well as in the honey bee [Zhou et al., 2015]. Unfortunately, further functional details on the nature of the traits affected by these losses cannot be inferred due to the lack of experimental results for such a large repertoire of genes with presumably complementary (epistatic) and overlapping (pleiotropic) effects. Still, this pattern constitutes another remarkable example of adaptive convergent gene loss, as a shift in environment (here, the social environment) may have rendered specific genes non-essential [Albalat and Cañestro, 2016]. This is reminiscent of studies investigating the transition from social to solitary bees [Wittwer et al., 2017] and provides valuable insight into the conditional importance of these genes under different social environments.

*Gr*s are instrumental for sensing nutrients [Freeman et al., 2014, Xu, 2020] and can be lost upon specialisation. For example, losses of *Gr*s and, to a much lesser extent of *Or* s, relative to their generalist sister species have been reported in two fly species which specialised on certain food resources [McBride and Arguello, 2007]. In ants, *Gr*s primarily respond to soluble tastants, and are thus presumably involved in foraging and harvesting [Zhou et al., 2012], though their exact functions still remain to be tested. As slave-maker ants outsource these tasks to host workers, thus likely rendering *Gr*s superfluous, we expected to observe random losses of *Gr*s in slave-maker ants across independent origins of social parasitism. Accordingly, we observed significant losses of *Gr*s in all slave-maker ants compared to their hosts.

While slave-maker ants lost a remarkable 50% of their *Gr* repertoire, no evidence was found for convergent losses of the exact same *Gr* orthologs in the three independently evolved parasites. Although there are much fewer *Gr* than *Or* orthologous clusters, we rule out that the lack of convergent loss in *Gr*s merely reflects a failure to detect convergent losses due to smaller sample size. Indeed, even if there were as many *Gr*s as *Or* s, our power analysis demonstrates that the probability to detect convergent losses in *Gr*s would still only be 35%, which is substantially less then the *>*90% found for *Or* s. Instead, these losses are consistent with strongly diminished need for gustatory reception through reliance on their host workers.

Host species in our study presented comparable numbers of *Or* s and *Gr*s in relation to non-host species. At first glance, this could indicate that parasitism does not lead to increased selective pressure for a more diverse chemosensory repertoire in host species. However, in host genomes, both *Or* s and *Gr*s underwent some degree of turnover, as losses and gains are both frequent and balance each other out (*Or* s: up to 8% of the repertoire was lost in host species, and a similar proportion underwent duplications; *Gr*s up to 8% of the repertoire was lost in hosts, and up to 30% underwent duplications; Figure 2 C and Figure 3 C). This underscores the potential of host species for rapid adaptive responses to changing environmental conditions [Simola et al., 2013]. Indeed, the evolutionary arms race for enemy recognition by the host and detection avoidance by the slave-maker ant species may select, through balancing or negative frequency-dependent selection, for a rapid turnover of nest-mate recognition substances and the associated receptors to detect them. Host species, in particular those in heavily parasitised locales, are thus under strong selection through destructive so-called “slave raids” to rapidly shift their chemosensory repertoire in order to maximise their chances of detecting an ever-changing parasite adversary [Jongepier et al., 2014, 2015, Scharf et al., 2011, Kleeberg et al., 2015, Jongepier and Foitzik, 2016]. The expansion or reduction of large clusters of tandem genes through unequal crossing over may have facilitated such a rapid turnover of receptors in host species.

In summary, our study shows that the slave-maker lifestyle in ants leads to a strong convergent reduction in *Or* and *Gr* genes and even convergent loss of specific *Or* orthologs. This study thus adds to the growing number of studies demonstrating the adaptiveness of gene loss associated with environmental or lifestyle change [Goldman-Huertas et al., 2015, Meyer et al., 2018]. While the results presented here provide compelling evidence for the adaptive nature of chemoreceptor losses in slave-maker ants during speciation, future studies will need to concentrate on the precise dynamics of the observed gene losses and the underlying genetic mechanisms. For example, it would be of interest to investigate the position of receptors lost in the slave-maker ants within genetic networks in the hosts, following the hypothesis that the loss of highly connected genes in parasites may have facilitated the emergence of a distinct phenotype [Helsen et al., 2020]. Additionally, studies of chemoreceptor gene expression would be relevant to investigate whether genes lost in one or several slave-maker ant species exhibit reduced expression in the remaining parasites, under the hypothesis that reduced gene expression precedes a gene loss.

## METHODS

### Sample collection

To generate genomic data across three independent origins of slave-making in ants, colonies of three slave-maker ant species (*Harpagoxenus sublaevis, Temnothorax ravouxi, Temnothorax americanus*), three host species (*Leptothorax acervorum, Temnothorax unifasciatus, Temnothorax longispinosus*) and two non-host outgroup species (*Temnothorax rugatulus, Temnothorax nylanderi*) were collected in the USA and Central Europe (Supplementary table S1). Multiple samples were pooled for each species to meet the requirements for whole-genome sequencing. For each species, two samples were prepared for WGS: 1) Pacbio Sequel long read library and 2) Illumina paired-end library. Selected host colonies were unparasitised, free-living colonies consisting of only a single species: the focal host. In the parasite colonies, all brood should in principle be parasite brood because host workers do not produce offspring. Nonetheless, to rule out that any of the samples taken from parasite colonies are actually host brood (*e.g*. relics from previous so-called “slave raids”), only pupae of castes that are morphologically distinct from their host were selected for sequencing (*i.e*. queen pupae for *Temnothorax ravouxi*, and queens and worker pupae for *Harpagoxenus sublaevis* and *Temnothorax americanus*). Depending on the availability of pupae for each species and the need to pool DNA extractions due to low DNA content, between 28 and 156 pupal samples were obtained for WGS.

### Genome sequencing

For each species, two samples were prepared for WGS: 1) PacBio Sequel long read library, aiming at 20 kb library construction and sequencing at 30*×* coverage; and 2) Illumina paired-end library with 150 bp long reads and 350 bp insert sizes. Only light coloured, unscleratized pupae were selected for sequencing because 1) the lack of a hard cuticle may reduce the risk of shearing high molecular weight DNA and 2) pupae shed most of their gut content which reduces contamination by gut bacteria. Pupae were sampled from their colony as they developed (*ca* 2 day intervals), snap frozen in liquid nitrogen and stored at -80^*°*^C. DNA extractions, quality checks, library preparation and sequencing were performed by Novogene under the umbrella of the Global Ant Genome Alliance [Boomsma et al., 2017]. For read statistics see Supplementary table S2.

### Genome assemblies and polishing

Several genome assembly strategies were explored and compared based on assembly size, contiguity and completeness (Supplementary tables S3-S10). For the final assembly, raw PacBio reads were assembled using the `Canu` [Koren et al., 2016] pipeline (parameter settings: correctedErrorRate=0.15), with K-mer based genome size estimates. The `Canu` assemblies was polished with `Pilon` (version 1.22; parameter settings: diploid, fix=all; Walker et al., 2014), using `Bowtie2`-aligned Illumina short reads (version 2.3.4.1; Langmead and Salzberg, 2012). Raw PacBio reads were mapped against the assemblies with `Minimap2` (version 2.1; settings: -ax map-pb; Li, 2018). Assemblies were then processed with `Purge Haplotigs` (Roach et al., 2018) and `FinisherSC` (in “fast” and “large” mode; Lam et al., 2015), followed by a final round of polishing with `Pilon`, `Arrow` (`VariantCaller` version 2.1.0) and again `Pilon`.

### Genome size estimates

Genome sizes were estimated using the following two strategies: 1) The K-mer distribution of the Illumina libraries, for which we used the `KmerCountExact` utility of `BBMap` [Bushnell, 2015]. This analysis was run for K-mer sizes ranging from 31 to 131, selecting the largest genome size estimate as input for the `CANU` [Koren et al., 2016] genome assemblies. 2) The coverage of the PacBio-based assembly, where we mapped the original PacBio reads back to the genome assembly using `Minimap2` (version 2.1; settings: `-ax map-pb`; Li, 2018) and determined the coverage frequency distribution with the `readhist` module of `Purge Haplotigs` [Roach et al., 2018]. The latter method is likely to yield higher and more correct genome size estimates than the former because large repetitive sequences are collapsed in the K-mer based estimate when they exceed Illumina read length. The coverage based genome size estimates of the final assemblies are very similar to the average genome size of Myrmicinae, which is 329.1 Mb [Tsutsui et al., 2008].

### Chemosensory receptor gene annotation

Reference protein predictions were obtained from the Ant Genomes Portal and functionally annotated based on their Pfam A domain content (*i.e*. gustatory receptors: “Trehalose recp” and “7tm 7”; odorant receptors: “7tm 6”; Finn et al., 2016; version 31). Chemoreceptor genes were manually annotated using a two-pass `tblastn/Exonerate` - `GeMoMa` - `WebApollo` workflow. In the first pass, these reference protein predictions were blasted against the assemblies using `tblastn` (version 2.6.0; e-value = 1 *×* 10^*-*3^Altschul et al., 1990). Annotations were then obtained with `Exonerate` (version 2.2.0; parameter settings: `--model protein2genome`; Slater and Birney, 2005), which was run on those genomic regions with a blast hit. In parallel, we obtained `GeMoMa` (version 1.4.2; Keilwagen et al., 2016) annotations, based on reference protein predictions as well as the RNA-seq libraries for intron predictions. `GeMoMa` annotations were filtered using `GAF`, either retaining only complete gene models or all predictions (*i.e*. parameter settings: `-r 0 -e 0`). Annotations were filtered based on their Pfam A annotation, retaining only those genes that had the defining domains. The `Exonerate`, `GeMoMa` (*i.e*. all predictions) and `GAF` gene models, as well as the mapped RNA-seq reads were used as evidence tracks for manual annotation using `WebApollo` (version 2.1.0; Lee et al., 2013), which involved manually leveraging all homology and expression evidence to construct the best gene model for each prediction. In the second pass, above workflow was repeated but now with all manual annotations from pass 1 as queries (*i.e*. from all focal species combined).

### Orthology clustering

To identify orthologous clusters, we took an explicit phylogenetic approach using the `Python` module `ETE3` (version 3.1.1; Huerta-Cepas et al. 2016). Specifically, multiple sequence alignments of odorant receptor (Or) and gustatory receptor (Gr) protein sequences from the eight focal species and nine reference species were obtained with `MAFFT` (version 7.310; Katoh and Standley 2013; parameter settings: `--maxiterate 1000 --localpair`). Gene trees were constructed with `Fasttree` (version 2.1; Price et al. 2010; parameter setting: `--pseudo`) and rooted with the Or co-receptor and the Trehalose receptors, for the *Or* and *Gr* tree respectively. The trees were traversed using `ETE3` and a clade was labelled as an orthologous cluster if a receptor from a reference species was found as outgroup. In total, 74% of the 379 orthologous clusters were 100% correctly identified and only 2.1% were completely missed. All clusters were thereafter manually curated and split or merged where necessary (for further details see Supplementary Section 3). Next, lowly supported branches (bootstrap value *<* 0.7) and species-specific expansions were collapsed.

### Convergent losses

For both chemosensory receptor families, we tested whether convergent loss occurred more often then expected by chance using a `binom.test` (R version 3.6.2; R Core Team 2019).

### Data availability statement

All raw DNA sequence data underlying this study as well as the novel genome assemblies will be deposited in the National Centre for Biotechnological Information (NCBI) Sequence Read Archive (SRA) and will be accessible upon publication of this manuscript (BioSample accession numbers pending). The odorant and gustatory receptor annotations will be made available in the Dryad Digital Repository (DOI pending) upon publication of this manuscript.

## Author contributions

The study was conceived by JH, EBB and SF, and was designed by EJ, JH, EBB and SF. EJ performed the experiments and analysed the data. BF, SF, JH, CG and AL contributed materials and analysis tools. EJ, AS, EBB, SF, JH and BF wrote the paper. Authors declare no conflict of interest.

## Acknowledgements

This work was supported by the Deutsche Forschungsgemeinschaft (Bo 2544/12-1, Fo 298/20, He 1623/40).

## Notes

### Competing Interest Statement

The authors have declared no competing interest.

## References

Ricard Albalat and Cristian Cañestro. Evolution by gene loss. Nature Reviews Genetics, 17(7):379, 2016. doi: https://doi.org/10.1038/nrg.2016.39.

Austin Alleman, Barbara Feldmeyer, and Susanne Foitzik. Comparative analyses of co-evolving host-parasite associations reveal unique gene expression patterns underlying slavemaker raiding and host defensive phenotypes. Scientific Reports, 8, 2018. doi: https://doi.org/10.1038/s41598-018-20262-y.

S. F. Altschul, W. Gish, W. Miller, E. W. Myers, and D. J. Lipman. Basic local alignment search tool. J. Mol. Biol., 215(3):403–410, Oct 1990. doi: https://doi.org/10.1016/S0022-2836(05)80360-2.

Laura Baxter, Sucheta Tripathy, Naveed Ishaque, Nico Boot, Adriana Cabral, Eric Kemen, Marco Thines, Audrey Ah-Fong, Ryan Anderson, Wole Badejoko, Peter Bittner-Eddy, Jeffrey L. Boore, Marcus C. Chibucos, Mary Coates, Paramvir Dehal, Kim Delehaunty, Suomeng Dong, Polly Downton, Berndard Dumas, Georgina Fabro, Catrina Fronick, Susan I. Fuerstenberg, Lucinda Fulton, Elodie Gaulin, Francine Govers, Linda Hughes, Sean Humphray, Rays H.Y. Jiang, Howard Judelson, Sophien Kamoun, Kim Kyung, Harold Meijer, Patrick Minx, Paul Morris, Joanne Nelson, Vipa Phuntumart, Dinah Qutob, Anne Rehmany, Alejandra Rougon-Cardoso, Peter Ryden, Trudy Torto-Alalibo, David Studholme, Yuanchao Wang, Joe Win, Jo Wood, Sandra W. Clifton, Jane Rogers, Guido van den Ackerveken, Jonathan D.G. Jones, John M. McDowel, Jim Beynon, and Brett M. Tyler. Signatures of adaptation to obligate biotrophy in the Hyaloperonospora arabidopsidis genome. Science, 330(6010):1549–1551, 2010. doi: 10.1126/science.1195203.

J. Beibl, R.J. Stuart, J. Heinze, and S. Foitzik. Six origins of slavery in formicoxenine ants. Insectes Sociaux, 52:291–297, 2005. doi: 10.1007/s00040-005-0808-y.

Calla Bernarda. Signatures of selection and evolutionary relevance of cytochrome p450s in plant-insect interactions. Current Opinion in Insect Science, 2020. doi: https://doi.org/10.1016/j.cois.2020.11.014.

Bonnie B Blaimer, Philip S Ward, Ted R Schultz, Brian L Fisher, and Seán G Brady. Paleotropical diversification dominates the evolution of the hyperdiverse ant tribe crematogastrini (hymenoptera: Formicidae). Insect Systematics and Diversity, 2(5):3, 2018. doi: https://doi.org/10.1093/isd/ixy013.

J. J. Boomsma, S. á. G. Brady, R. R. Dunn, J. Gadau, J Heinze, L. Keller, N. J. Sanders, L. Schrader, T. R. Schultz, Sundstöm L., P. S. Ward, Wcislo W. T., Zhang G., and The GAGA Consortium. The global ant genomics alliance (gaga). Myrmecological News, 25:61–66, 2017.

A. Buschinger. Evolution of social parasitism in ants. Trends in Ecology and Evolution, 1:155–160, 1986. doi: https://doi.org/10.1016/0169-5347(86)90044-3.

Brian Bushnell. Bbmap short read aligner, and other bioinformatics tools. http://sourceforge.net/projects/bbmap/, 2015.

Charles Darwin. On the Origin of Species by Means of Natural Selection. John Murray, London, 1859.

Guy Drouin, Jean-Rémi Godin, and Benoît Pagé. The genetics of vitamin c loss in vertebrates. Current Genomics, 12(5):371–378, 2011. doi: https://doi.org/10.2174/138920211796429736.

C. Emery. über den ursprung der dulotischen, parasitischen und myrmekophilen ameisen. Biologisches Centralblatt, 29:352–362, 1909.

Patamarerk Engsontia, Unitsa Sangket, Hugh M. Robertson, and Chutamas Satasook. Diversification of the ant odorant receptor gene family and positive selection on candidate cuticular hydrocarbon receptors. BMC Research Notes, 8, 2015. doi: https://doi.org/10.1186/s13104-015-1371-x.

B. Feldmeyer, D. Elsner, A. Alleman, and S. Foitzik. Species-specific genes under selection characterize the co-evolution of slavemaker and host lifestyles. BMC Evolutionary Biology, 17(1):237, Dec 2017. ISSN 1471-2148. doi: 10.1186/s12862-017-1078-9”.

R. D. Finn, P. Coggill, R. Y. Eberhardt, S. R. Eddy, J. Mistry, A. L. Mitchell, S. C. Potter, M. Punta, M. Qureshi, A. Sangrador-Vegas, G. A. Salazar, J. Tate, and A. Bateman. The Pfam protein families database: towards a more sustainable future. Nucleic Acids Res., 44(D1):D279–285, Jan 2016. doi: https://doi.org/10.1093/nar/gkv1344.

Susanne Foitzik, Christopher J DeHeer, Daniel N Hunjan, and Joan M Herbers. Coevolution in host–parasite systems: behavioural strategies of slave–making ants and their hosts. Proceedings of the Royal Society of London. Series B: Biological Sciences, 268(1472):1139–1146, 2001. doi: https://doi.org/10.1098/rspb.2001.1627.

Erica Gene Freeman, Zev Wisotsky, and Anupama Dahanukar. Detection of sweet tastants by a conserved group of insect gustatory receptors. Proceedings of the National Academy of Sciences, 111(4):1598–1603, 2014. doi: https://doi.org/10.1073/pnas.1311724111.

Benjamin Goldman-Huertas, Robert F Mitchell, Richard T Lapoint, Cécile P Faucher, John G Hildebrand, and Noah K Whiteman. Evolution of herbivory in drosophilidae linked to loss of behaviors, antennal responses, odorant receptors, and ancestral diet. Proceedings of the National Academy of Sciences, 112 (10):3026–3031, 2015. doi: https://doi.org/10.1073/pnas.1424656112.

Cristina Guijarro-Clarke, Peter WH Holland, and Jordi Paps. Widespread patterns of gene loss in the evolution of the animal kingdom. Nature Ecology & Evolution, 4(4):519–523, 2020. doi: https://doi.org/10.1038/s41559-020-1129-2.

J Heinze. The reproductive potential of workers in slave-making ants. Insectes sociaux, 43(3):319–328, 1996.

Jana Helsen, Karin Voordeckers, Laura Vanderwaeren, Toon Santermans, Maria Tsontaki, Kevin J Verstrepen, and Rob Jelier. Gene loss predictably drives evolutionary adaptation. Molecular Biology and Evolution, 2020. doi: https://doi.org/10.1093/molbev/msaa172.

B. Hölldobler and E. O. Wilson. The Ants. Harvard University Press, Cambridge, Massachusetts, 1990.

Bert Hölldobler. The chemistry of social regulation: multicomponent signals in ant societies. Proceedings of the National Academy of Sciences, 92(1):19–22, 1995.

Jaime Huerta-Cepas, François Serra, and Peer Bork. ETE 3: Reconstruction, Analysis, and Visualization of Phylogenomic Data. Molecular Biology and Evolution, 33(6):1635–1638, 02 2016. ISSN 0737-4038. doi: 10.1093/molbev/msw046.

Michelle T Hulin, Andrew D Armitage, Joana G Vicente, Eric B Holub, Laura Baxter, Helen J Bates, John W Mansfield, Robert W Jackson, and Richard J Harrison. Comparative genomics of Pseudomonas syringae reveals convergent gene gain and loss associated with specialization onto cherry (prunus avium). New Phytologist, 219(2):672–696, 2018. doi: https://doi.org/10.1111/nph.15182.

E Jongepier, I Kleeberg, and S Foitzik. The ecological success of a social parasite increases with manipulation of collective host behaviour. Journal of evolutionary biology, 28(12):2152–2162, 2015. doi: https://doi.org/10.1111/jeb.12738.

Evelien Jongepier and Susanne Foitzik. Ant recognition cue diversity is higher in the presence of slavemaker ants. Behavioral Ecology, 27(1):304–311, 2016. doi: https://doi.org/10.1093/beheco/arv153.

Evelien Jongepier, Isabelle Kleeberg, Sylwester Job, and Susanne Foitzik. Collective defence portfolios of ant hosts shift with social parasite pressure. Proceedings of the Royal Society B: Biological Sciences, 281 (1791):20140225, 2014. doi: https://doi.org/10.1098/rspb.2014.0225.

Kazutaka Katoh and Daron M. Standley. MAFFT Multiple Sequence Alignment Software Version 7: Improvements in Performance and Usability. Molecular Biology and Evolution, 30(4):772–780, 01 2013. ISSN 0737-4038. doi: 10.1093/molbev/mst010.

Jens Keilwagen, Michael Wenk, Jessica L. Erickson, Martin H. Schattat, Jan Grau, and Frank Hartung. Using intron position conservation for homology-based gene prediction. Nucleic Acids Research, 44(9): e89, 2016. doi: 10.1093/nar/gkw092.

Ewen F Kirkness, Brian J Haas, Weilin Sun, Henk R Braig, M Alejandra Perotti, John M Clark, Si Hyeock Lee, Hugh M Robertson, Ryan C Kennedy, Eran Elhaik, et al. Genome sequences of the human body louse and its primary endosymbiont provide insights into the permanent parasitic lifestyle. Proceedings of the National Academy of Sciences, 107(27):12168–12173, 2010. doi: https://doi.org/10.1073/pnas.1003379107.

Isabelle Kleeberg, Evelien Jongepier, Sylwester Job, and Susanne Foitzik. Geographic variation in social parasite pressure predicts intraspecific but not interspecific aggressive responses in hosts of a slavemaking ant. Ethology, 121(7):694–702, 2015. doi: https://doi.org/10.1111/eth.12384.

Fyodor A Kondrashov. Gene duplication as a mechanism of genomic adaptation to a changing environment. Proceedings of the Royal Society B: Biological Sciences, 279(1749):5048–5057, 2012. doi: https://doi.org/10.1098/rspb.2012.1108.

S Koren, B. P. Walenz, K. Berlin, J. R. Miller, and A. M. Phillippy. Canu: scalable and accurate long-read assembly via adaptive k-mer weighting and repeat separation. bioRxiv, August 2016.

Ka-Kit Lam, Kurt LaButti, Asif Khalak, and David Tse. Finishersc: a repeat-aware tool for upgrading de novo assembly using long reads. Bioinformatics, 31(19):3207–3209, 2015. doi: 10.1093/bioinformatics/btv280.

Ben Langmead and Steven L. Salzberg. Fast gapped-read alignment with bowtie 2. Nature Methods, 9: 357–359, 2012. doi: 10.1038/nmeth.1923.

Eduardo Lee, Gregg A. Helt, Justin T. Reese, Monica C. Munoz-Torres, Chris P. Childers, Robert M. Buels, Lincoln Stein, Ian H. Holmes, Christine G. Elsik, and Suzanna E. Lewis. Web apollo: a web-based genomic annotation editing platform. Genome Biology, 14(8):R93, Aug 2013. doi: 10.1186/gb-2013-14-8-r93.

Heng Li. Minimap2: pairwise alignment for nucleotide sequences. Bioinformatics, 34(18):3094–3100, 2018. doi: 10.1093/bioinformatics/bty191.

Carolyn S McBride and J Roman Arguello. Five drosophila genomes reveal nonneutral evolution and the signature of host specialization in the chemoreceptor superfamily. Genetics, 177(3):1395–1416, 2007. doi: https://doi.org/10.1534/genetics.107.078683.

Sean K McKenzie and Daniel JC Kronauer. The genomic architecture and molecular evolution of ant odorant receptors. Genome research, 28(11):1757–1765, 2018. doi: 10.1101/gr.237123.118.

Sean K. McKenzie, Ingrid Fetter-Pruneda, Vanessa Ruta, and Daniel J. C. Kronauer. Transcriptomics and neuroanatomy of the clonal raider ant implicate an expanded clade of odorant receptors in chemical communication. Proceedings of the National Academy of Sciences, 113(49):14091–14096, 2016. doi: 10.1073/pnas.1610800113.

Wynn K Meyer, Jerrica Jamison, Rebecca Richter, Stacy E Woods, Raghavendran Partha, Amanda Kowalczyk, Charles Kronk, Maria Chikina, Robert K Bonde, Daniel E Crocker, Joseph Gaspard, Janet M Lanyon, Judit Marsillach, Clement E Furlong, and Nathan L Clark. Ancient convergent losses of paraoxonase 1 yield potential risks for modern marine mammals. Science, 361(6402):591–594, 2018. doi: 10.1126/science.aap7714.

Makedonka Mitreva, Douglas P Jasmer, Dante S Zarlenga, Zhengyuan Wang, Sahar Abubucker, John Martin, Christina M Taylor, Yong Yin, Lucinda Fulton, Pat Minx, Shiaw-Pyng Yang, Wesley C Warren, Robert S Fulton, Veena Bhonagiri, Xu Zhang, Kym Hallsworth-Pepin, Sandra W Clifton, James P McCarter, Judith Appleton, Elaine R Mardis, and Richard K Wilson. The draft genome of the parasitic nematode trichinella spiralis. Nature genetics, 43(3):228–235, 2011. doi: https://doi.org/10.1038/ng.769.

Maynard V Olson. When less is more: gene loss as an engine of evolutionary change. The American Journal of Human Genetics, 64(1):18–23, 1999.

Gregory M Pask, Jesse D Slone, Jocelyn G Millar, Prithwiraj Das, Jardel A Moreira, Xiaofan Zhou, Jan Bello, Shelley L Berger, Roberto Bonasio, Claude Desplan, Danny Reinberg, Jürgen Liebig, Laurence J Zwiebel, and Anandasankar Ray. Specialized odorant receptors in social insects that detect cuticular hydrocarbon cues and candidate pheromones. Nature communications, 8(1):1–11, 2017. doi: https://doi.org/10.1038/s41467-017-00099-1.

Robert Poulin and Haseeb S Randhawa. Evolution of parasitism along convergent lines: from ecology to genomics. Parasitology, 142(S1):S6–S15, 2015. doi: https://doi.org/10.1017/S0031182013001674.

Matthew Prebus. Insights into the evolution, biogeography and natural history of the acorn ants, genus temnothorax mayr (hymenoptera: Formicidae). BMC Evolutionary Biology, 17(1):250, Dec 2017. ISSN 1471-2148. doi: 10.1186/s12862-017-1095-8.

Morgan N. Price, Paramvir S. Dehal, and Adam P. Arkin. Fasttree 2 – approximately maximum-likelihood trees for large alignments. PLOS ONE, 5(3):1–10, 03 2010. doi: 10.1371/journal.pone.0009490.

Wenfeng Qian and Jianzhi Zhang. Genomic evidence for adaptation by gene duplication. Genome research, 24(8):1356–1362, 2014. doi: 10.1101/gr.172098.114.

R Core Team. R: A Language and Environment for Statistical Computing. R Foundation for Statistical Computing, Vienna, Austria, 2019. URL https://www.R-project.org/.

FE Regnier and EO Wilson. Chemical communication and” propaganda” in slave-maker ants. Science, 172 (3980):267–269, 1971.

Michael J Roach, Simon A Schmidt, and Anthony R Borneman. Purge haplotigs: Synteny reduction for third-gen diploid genome assemblies. bioRxiv, 2018. doi: 10.1101/286252.

Inon Scharf, Tobias Pamminger, and Susanne Foitzik. Differential response of ant colonies to intruders: attack strategies correlate with potential threat. Ethology, 117(8):731–739, 2011. doi: https://doi.org/10.1111/j.1439-0310.2011.01926.x.

Daniel F Simola, Lothar Wissler, Greg Donahue, Robert M Waterhouse, Martin Helmkampf, Julien Roux, Sanne Nygaard, Karl M Glastad, Darren E Hagen, Lumi Viljakainen, et al. Social insect genomes exhibit dramatic evolution in gene composition and regulation while preserving regulatory features linked to sociality. Genome research, 23(8):1235–1247, 2013. doi: 10.1101/gr.155408.113.

Guy St C Slater and Ewan Birney. Automated generation of heuristics for biological sequence comparison. BMC bioinformatics, 6:31, 2005. doi: 10.1186/1471-2105-6-31.

Jesse D Slone, Gregory M Pask, Stephen T Ferguson, Jocelyn G Millar, Shelley L Berger, Danny Reinberg, Jürgen Liebig, Anandasankar Ray, and Laurence J Zwiebel. Functional characterization of odorant receptors in the ponerine ant, Harpegnathos saltator. Proceedings of the National Academy of Sciences, 114 (32):8586–8591, 2017. doi: https://doi.org/10.1073/pnas.1704647114.

Evgeni V Sokurenko, David L Hasty, and Daniel E Dykhuizen. Pathoadaptive mutations: gene loss and variation in bacterial pathogens. Trends in microbiology, 7(5):191–195, 1999. doi: https://doi.org/10.1016/S0966-842X(99)01493-6.

Guiling Sun, Yuxing Xu, Hui Liu, Ting Sun, Jingxiong Zhang, Christian Hettenhausen, Guojing Shen, Jinfeng Qi, Yan Qin, Jing Li, Lei Wang, Wei Chang, Guo Zhenhua, Ian T Baldwin, and Jianqiang Wu. Large-scale gene losses underlie the genome evolution of parasitic plant cuscuta australis. Nature communications, 9(1):1–8, 2018. doi: https://doi.org/10.1038/s41467-018-04721-8.

Neil D. Tsutsui, Andrew V. Suarez, Joseph C. Spagna, and J. Spencer Johnston. The evolution of genome size in ants. BMC Evolutionary Biology, 8(1):64, 2008. ISSN 1471-2148. doi: 10.1186/1471-2148-8-64.

Bruce J. Walker, Thomas Abeel, Terrance Shea, Margaret Priest, Amr Abouelliel, Sharadha Sakthikumar, Christina A. Cuomo, Qiandong Zeng, Jennifer Wortman, Sarah K. Young, and Ashlee M. Earl. Pilon: An integrated tool for comprehensive microbial variant detection and genome assembly improvement. PLOS ONE, 9(11):1–14, 11 2014. doi: 10.1371/journal.pone.0112963.

Philip S Ward. The phylogeny and evolution of ants. Annual Review of Ecology, Evolution, and Systematics, 45:23–43, 2014. doi: https://doi.org/10.1146/annurev-ecolsys-120213-091824.

Brooke L Whitelaw, Ira R Cooke, Julian Finn, Rute R da Fonseca, Elena A Ritschard, MTP Gilbert, Oleg Simakov, and Jan M Strugnell. Adaptive venom evolution and toxicity in octopods is driven by extensive novel gene formation, expansion, and loss. GigaScience, 9(11):giaa120, 2020. doi: https://doi.org/10.1093/gigascience/giaa120.

Bernadette Wittwer, Abraham Hefetz, Tovit Simon, Li EK Murphy, Mark A Elgar, Naomi E Pierce, and Sarah D Kocher. Solitary bees reduce investment in communication compared with their social relatives. Proceedings of the National Academy of Sciences, 114(25):6569–6574, 2017. doi: https://doi.org/10.1073/pnas.1620780114.

Wei Xu. How do moth and butterfly taste?—molecular basis of gustatory receptors in lepidoptera. Insect science, 27(6):1148–1157, 2020. doi: https://doi.org/10.1111/1744-7917.12718.

Xiaofan Zhou, Jesse D Slone, Antonis Rokas, Shelley L Berger, Jürgen Liebig, Anandasankar Ray, Danny Reinberg, and Laurence J Zwiebel. Phylogenetic and transcriptomic analysis of chemosensory receptors in a pair of divergent ant species reveals sex-specific signatures of odor coding. PLoS Genet, 8(8):e1002930, 2012. doi: https://doi.org/10.1371/journal.pgen.1002930.

Xiaofan Zhou, Antonis Rokas, Shelley L. Berger, Jürgen Liebig, Anandasankar Ray, and Laurence J. Zwiebel. Chemoreceptor evolution in Hymenoptera and its implications for the evolution of eusociality. Genome Biology and Evolution, 7(8):2407–2416, 08 2015. ISSN 1759-6653. doi: 10.1093/gbe/evv149.

Yanyan Zhou, Xiaotong Li, Susumu Katsuma, Yusong Xu, Liangen Shi, Toru Shimada, and Huabing Wang. Duplication and diversification of trehalase confers evolutionary advantages on lepidopteran insects. Molecular Ecology, 28(24):5282–5298, 2019. doi: https://doi.org/10.1111/mec.15291.

