## Supplementary for "Convergent loss of chemoreceptors across independent origins of slave-making in ants"

### SUPPLEMENTARY MATERIAL

Evelien Jongepier[a,b,*], Alice Séguret[a], Anton Labutin[a], Barbara Feldmeyer[c], Claudia Gstöttl[d], Susanne Foitzik[e,*], Jürgen Heinze[d], and Erich Bornberg-Bauer[a,*]

[a]Institute for Evolution and Biodiversity, Westfälische Wilhelms University, Münster, Germany

[b]Institute for Biodiversity and Ecosystem Dynamics, University of Amsterdam, Amsterdam, The Netherlands

[c]Senckenberg Biodiversity and Climate Research Centre, Frankfurt am Main, Germany

[d]Institute for Zoology, University of Regensburg, Regensburg, Germany

[e]Institute of Organismic and Molecular Evolution, Johannes Gutenberg University, Mainz, Germany

2021-05-11

### SUPPLEMENTARY MATERIAL

#### 1 Genome assemblies

Several genome assembly strategies were explored and compared based on assembly size (section Genome size estimates), contiguity (assessed with `QUAST`; version 3.1; Gurevich et al., 2013) and completeness (assessed with `BUSCO`; version 3.0.2; insect db; Simão et al., 2015). Of particular concern was that several assembly sizes were (much) larger than expected (see section Genome size estimates) and these assemblies contained a high number of duplicated orthologs which usually occur as single-copy genes in insect genomes (*i.e.* duplicated `BUSCOs`). This assembly inflation is likely due to the fact that several individuals had to be pooled in order to obtain enough high molecular weight DNA for these tiny ants (section Sample collection). Thus, the samples used for PacBio library prep were polyploid, which can result in the same polymorphic genomic region being represented in the assembly as separate contigs, rather than as a contig and an associated haplotig. The main aim of the following assembly strategies was thus to reduce redundancy and obtain a haploid genome assembly for each of our focal species.

1. `MaSuRCA`: For most of our focal species, a preliminary assembly was generated with `MaSuRCA` (version 3.2.6; default parameter settings; Zimin et al., 2013) based on the raw, untrimmed Illumina reads only. This assembly served to error correct the PacBio reads (see 2).

2. `HALC-CANU`: Raw PacBio reads were error corrected prior to assembly with the help of the preliminary `MaSuRCA` assembly. The PacBio reads were then aligned to the `MaSuRCA` assembly using `Blasr` (`smrtanalyses` version 2.3.0; parameter settings: maxScore=2000, minMatch=8, nCandidates=30, bestn=20, m=5; Chaisson and Tesler, 2012). Aligned reads were error-corrected with `HALC` (date: 04.2018; default parameter settings of the `HALC` wrapper script; Bao and Lan, 2017), making use of LorDEC (version 0.7; Salmela and Rivals, 2014). The resulting corrected, trimmed and split PacBio reads were then assembled using `CANU` (version 1.7; Koren et al., 2016; default parameter settings and the K-mer based genome size estimates, see section Genome size estimates).

3. $\texttt{CANU}_{\texttt{default}}$: Raw PacBio reads were error corrected, trimmed, split and assembled using the `CANU` [Koren et al., 2016] pipeline with default parameter settings and the K-mer based genome size estimates (see section Genome size estimates).

4. CANU$_{\text{high error}}$: Raw PacBio reads were error corrected, trimmed, split and assembled using the CANU [Koren et al., 2016] pipeline (using K-mer based genome size estimates, see section Genome size estimates). Instead of using default parameter settings as in 2, we tweaked the CANU error rate and coverage settings in order to smash haplotypes together (`correctedErrorRate=0.15, corOutCoverage=200`; see also Canu FAQ (here)).

Not all assembly methods were applied to all species: assemblies were skipped if we had already decided to proceed with an alternative approach based on results in the other species (indicated by empty columns in Supplementary tables S3-S10).

#### 2 Genome polishing

The Canu assemblies were polished with Pilon (version 1.22; parameter settings: diploid, fix=all; Walker et al., 2014), using the Illumina short reads, which were first aligned to the Canu assembly using Bowtie2 (version 2.3.4.1; Langmead and Salzberg, 2012).

The $\text{CANU}_{\text{high error}}$ assemblies was further processed with Purge Haplotigs (Roach et al., 2018) to identify syntenic pairs of contigs and discard one of them. Hereto, the raw PacBio reads were mapped against the Canu genome assembly with Minimap2 (version 2.1; settings: -ax map-pb; Li, 2018). Assessment of genome read coverage and read depth cutoffs, as well as the actual purging of haplotigs was done with the Purge Haplotigs pipeline [Roach et al., 2018].

Coverage frequency distributions provide insight into ploidy: In a haploid genome assembly where each genomic location is represented only once, the coverage frequency distribution is unimodal and symmetrical around its mode. If the same genomic location is erroneously assembled twice due to heterozygosity in diploid samples, then each of these contigs will only get half of the coverage, causing a second mode at half the coverage of the haploid peak. Given that we had polyploid rather than diploid source material, no bimodality in the read depth frequency distributions could be observed. Instead, an over-representation of low coverages was observed in the $\text{CANU}_{\text{default}}$, and to a lesser extent in $\text{CANU}_{\text{high error}}$, but not in the final Purge Haplotigs polished assemblies. Importantly, purging haplotigs did not affect BUSCO scores, with the exception of a substantial drop in duplicated BUSCOs (Tables S6-S10).

The Purge Haplotigs-polished assemblies were then upgraded with FinisherSC (in "fast" and "large" mode; Lam et al., 2015), followed by a final round of polishing with Arrow (VariantCaller version 2.1.0) and Pilon (version 1.22; parameter settings: diploid, fix=all; Walker et al., 2014). Arrow can correct large structural errors as it is based on the raw PacBio reads. Quiver aims to correct small errors based on the short, but high quality Illumina reads. To run Arrow, raw PacBio reads were aligned to the assembly with Minimap2 and converted to a PacBio-compatible bam using pbbamify (version 0.13.2).

#### 3 Orthology and Phylogeny

Preliminary analyses using `OrthoMCL` or `OrthoFinder` yielded very poor orthology clustering results. This is probably due to the large number of paralogs, and the close relatedness among our focal species. Visual inspection of the Or and Gr protein trees however showed that clear orthologous clusters could be identified, especially when the receptors of more distantly related outgroup species were considered. We therefore chose to use an explicit phylogenetic approach to identify orthologous clusters using the `Python` module `ETE3` (version 3.1.1; Huerta-Cepas et al. 2016). Specifically, multiple sequence alignments of Or and Gr protein sequences from the eight focal species and nine reference species (using reference genomes available on the Ant Genomes Portal) were obtained with `MAFFT` (version 7.310; Katoh and Standley 2013; parameter settings: `--maxiterate 1000 --localpair`). Gene trees were constructed with `Fasttree` (version 2.1; Price et al. 2010; parameter setting: `--pseudo`) and rooted based on the Or co-receptor and the Trehalose receptors, for the *Or* and *Gr* tree respectively. The trees were traversed using `ETE3` and a clade was labelled as an orthologous cluster if a receptor from a reference species was found as outgroup.

Obtaining a single multiple sequence alignment for all 4 759 *Or*s present in the nine reference and eight focal species required more than the available 756 GB of RAM. Therefore, (i) the Or protein tree of the eight focal species was split into four sub-trees. (ii) The Or protein sequences of the reference species were `BLASTP`-ed against the Or protein sequences of the focal species. (iii) Each reference species Or was assigned to the same sub-tree as its best blast hit, yielding four separate sets of Or protein sequences from all 17 species. Multiple sequence alignment, phylogenetic tree construction and orthology assignment was then done as described above for each of the four sub-trees separately.

Visual inspection confirmed correct automated identification of 58.3% the 72 final orthologous Gr clusters and 77.5% of the 307 orthologous Or clusters. Manual curation resulted in the merging of five Gr clusters, the identification of an additional four clusters, and the splitting of eleven initial clusters in two or more orthologous groups based on the presence of *H. sublaevis* and/or *L. acervorum* as an outgroup to the other focal species. For the Ors, manual curation resulted in 25 mergings, six splits and the identification of four additional orthologous clusters.

Table S1: Sample information.

| Species | Colony ID | Latitude | Longitude | Collection date | Sample |
|---|---|---|---|---|---|
| *Temnothorax unifasciatus* | Tuni_6 | 49°09'51.2" | 11°56'49.7" | 06-10-2016 | 112 pupae |
| *Temnothorax ravouxi* | Mrav_6 | 49°00'39.0" | 11°57'12.0" | 21-06-2018 | 156 queen pupae |
| *Harpagoxenus sublaevis* | Hsub_1 | 49°20'40.4" | 11°15'37.4" | 10-05-2017 | 28 female pupae |
| *Leptothorax acervorum* | Lace_23 | 49°21'42.0" | 11°17'17.0" | 10-05-2017 | 75 pupae |
| *Temnothorax americanus* | Tame_NY17_H226 | 42°31'36.9" | -74°10'15.6" | 25-05-2017 | 42 female pupae |
| *Temnothorax longispinosus* | Tlon_NY17_J1612 | 42°31'36.9" | -74°10'15.6" | 25-05-2017 | 136 pupae |
| *Temnothorax rugatulus* | Trug_M654 | 31°51'1.8" | -109°19'32.2" | 20-08-2015 | 140 pupae |
| *Temnothorax nylanderi* | Tnyl_W156 | 49°48'38.0" | 7°52'07.9" | 15-08-2016 | 148 pupae |

Table S2: Genomic read statistics.

| | Illumina reads | | PacBio reads | | | |
|---|---|---|---|---|---|---|
| Species | Gbases | No. read pairs | Gbases | No. reads | N50 | N90 |
| *Temnothorax unifasciatus* | 16.636 | 55 454 527 | 14.647 | 2 115 430 | 10 295 | 4 256 |
| *Temnothorax ravouxi* | 16.358 | 54 527 419 | 24.727 | 2 810 494 | 11 186 | 5 900 |
| *Harpagoxenus sublaevis* | 18.076 | 60 254 049 | 16.094 | 2 758 201 | 8 940 | 3 423 |
| *Leptothorax acervorum* | 15.666 | 52 218 260 | 16.353 | 2 077 285 | 12 114 | 4 805 |
| *Temnothorax americanus* | 11.725 | 38 825 160 | 13.919 | 1 721 559 | 10 692 | 5 280 |
| *Temnothorax longispinosus* | 28.691 | 95 002 424 | 19.476 | 2 579 317 | 10 916 | 4 465 |
| *Temnothorax rugatulus* | 15.664 | 52 214 755 | 18.610 | 2 282 653 | 11 733 | 4 900 |
| *Temnothorax nylanderi* | 18.267 | 60 889 461 | 17.148 | 2 002 476 | 12 477 | 5 111 |

Table S3: *Harpagoxenus sublaevis* assembly statistics. The column `MaSuRCA` contains the statistics for the assembly generated with `MaSuRCA`, the column `HALC-Canu`$_{\texttt{def}}$ contains the statistics for the assembly generated using `CANU` with default parameter settings, using reads error-corrected with `HALC`, the column `Canu`$_{\texttt{def}}$`-Pilon` contains statistics for the assembly generated using `CANU` with default parameter settings, followed by a round of polishing with `Pilon`, and the column `Canu`$_{\texttt{err}}$`-Pilon` indicates statistics for the assembly generated using `CANU` with high error rates, followed by a round of polishing with `Pilon`. The `Canu`$_{\texttt{err}}$ assemblies were further processed with `Purge Haplotigs` (statistics presented in the column `PurgeHaplotigs`), then upgraded with `FinisherSC` (statistics presented in column `FinisherSC`) and a final round of polishing with `Arrow` and `Pilon` (statistics presented in the last column). Not all assembly methods were applied to all species (hence the empty columns) because this was not necessary to evaluate the performance of the applied methods. Only the assembly presented in the last column was used for further analyses.

| Assembly | `MaSuRCA` | `HALC-Canu`$_{\texttt{def}}$ | `Canu`$_{\texttt{def}}$`-Pilon` | `Canu`$_{\texttt{err}}$`-Pilon` | `PurgeHaplotigs` | `FinisherSC` | `Arrow-Pilon` |
|---|---|---|---|---|---|---|---|
| No. contigs | 164 918 | - | 7 355 | 5 956 | 2 794 | 2 750 | 2 750 |
| Total length | 376 588 678 | - | 429 220 703 | 392 891 886 | 341 470 629 | 341 284 157 | 341 523 168 |
| Largest contig | 87 368 | - | 6 171 555 | 4 362 322 | 4 362 322 | 4 360 839 | 4 363 893 |
| GC (%) | 39.10 | - | 39.15 | 39.17 | 39.00 | 39.00 | 39.00 |
| N50 | 5 179 | - | 224 287 | 260 672 | 390 745 | 390 729 | 391 101 |
| N75 | 1 925 | - | 42 041 | 54 059 | 94 152 | 94 153 | 94 448 |
| L50 | 18 273 | - | 327 | 258 | 176 | 176 | 176 |
| L75 | 47 420 | - | 1 503 | 1 195 | 651 | 650 | 649 |
| Complete BUSCO's (%) | 87.8 | - | 98.1 | 98.2 | 98.1 | 97.5 | 98.1 |
| Complete, single copy BUSCO's (%) | 86.3 | - | 86.9 | 94.3 | 96.7 | 96.3 | 96.7 |
| Complete, duplicated BUSCO's (%) | 1.5 | - | 11.2 | 3.9 | 1.4 | 1.2 | 1.4 |
| Fragmented BUSCO's (%) | 8.5 | - | 0.7 | 0.7 | 0.7 | 1.2 | 0.7 |
| Missing BUSCO's (%) | 3.7 | - | 1.2 | 1.1 | 1.2 | 1.3 | 1.2 |

Table S4: *Leptothorax acervorum* assembly statistics. For further details, see Table S3 legend.

| Assembly | MaSuRCA | HALC-Canu$_{def}$ | Canu$_{def}$-Pilon | Canu$_{err}$-Pilon | PurgeHaplotigs | FinisherSC | Arrow-Pilon |
|---|---|---|---|---|---|---|---|
| No. contigs | 79 589 | 10 676 | 7 544 | 5 433 | 1 545 | 1 269 | 1 269 |
| Total length | 348 003 143 | 411 018 547 | 516 250 221 | 455 819 920 | 346 747 927 | 345 140 810 | 344 708 930 |
| Largest contig | 382 012 | 774 699 | 4 002 204 | 6 338 506 | 6 338 506 | 6 338 506 | 6 338 531 |
| GC (%) | 38.57 | 38.51 | 38.58 | 38.66 | 38.68 | 38.67 | 38.67 |
| N50 | 21 005 | 67 835 | 211 559 | 298 336 | 605 042 | 729 587 | 730 108 |
| N75 | 4 656 | 27 793 | 43 009 | 65 476 | 194 900 | 247 721 | 248 295 |
| L50 | 3 332 | 1 301 | 458 | 247 | 119 | 104 | 104 |
| L75 | 12 722 | 3 865 | 1 990 | 1 140 | 380 | 310 | 310 |
| Complete BUSCO's (%) | 95.3 | - | 98.2 | 98.3 | 98.1 | 98.0 | 98.3 |
| Complete, single copy BUSCO's (%) | 94.5 | - | 80.9 | 92.1 | 96.0 | 96.1 | 96.7 |
| Complete, duplicated BUSCO's (%) | 0.8 | - | 17.3 | 6.2 | 2.1 | 1.9 | 1.6 |
| Fragmented BUSCO's (%) | 2.8 | - | 0.8 | 0.7 | 0.7 | 0.7 | 0.6 |
| Missing BUSCO's (%) | 1.9 | - | 1.0 | 1.0 | 1.2 | 1.3 | 1.1 |



Table S5: *Temnothorax ravouxi* assembly statistics. For further details, see Table S3 legend.

| Assembly | MaSuRCA | HALC-Canu$_{def}$ | Canu$_{def}$-Pilon | Canu$_{err}$-Pilon | PurgeHaplotigs | FinisherSC | Arrow-Pilon |
|---|---|---|---|---|---|---|---|
| No contigs | - | - | - | 18 188 | 5 018 | 4 380 | 4 380 |
| Total length | - | - | - | 596 230 198 | 366 250 529 | 362 215 514 | 362 408 415 |
| Largest contig | - | - | - | 1 215 712 | 1 216 519 | 1 285 007 | 1 282 434 |
| GC (%) | - | - | - | 38.67 | 38.69 | 38.69 | 38.70 |
| N50 | - | - | - | 51 549 | 106 656 | 128 643 | 128 698 |
| N75 | - | - | - | 22 875 | 56 975 | 66 149 | 66 202 |
| L50 | - | - | - | 2 477 | 901 | 751 | 751 |
| L75 | - | - | - | 7 007 | 2 087 | 1 733 | 1 733 |
| Complete BUSCO's (%) | - | - | - | - | - | - | 95.7 |
| Complete, single copy BUSCO's (%) | - | - | - | - | - | - | 88.0 |
| Complete, duplicated BUSCO's (%) | - | - | - | - | - | - | 7.7 |
| Fragmented BUSCO's (%) | - | - | - | - | - | - | 1.2 |
| Missing BUSCO's (%) | - | - | - | - | - | - | 3.1 |

Table S6: *Temnothorax unifasciatus* assembly statistics. For further details, see Table S3 legend.

| Assembly | MaSuRCA | HALC-Canu$_{def}$ | Canu$_{def}$-Pilon | Canu$_{err}$-Pilon | PurgeHaplotigs | FinisherSC | Arrow-Pilon |
|---|---|---|---|---|---|---|---|
| No. contigs | 47 056 | 12 586 | - | 5 915 | 2 964 | 2 496 | 2 496 |
| Total length | 294 647 559 | 464 253 540 | - | 365 527 575 | 317 439 015 | 314 940 905 | 314 490 334 |
| Largest contig | 347 969 | 1 205 371 | - | 2 812 578 | 2 812 578 | 2 906 488 | 2 901 263 |
| GC (%) | 39.20 | 38.67 | - | 38.72 | 38.80 | 38.80 | 38.81 |
| N50 | 25 189 | 66 013 | - | 214 554 | 290 891 | 360 251 | 359 731 |
| N75 | 8 152 | 25 538 | - | 48 796 | 92 794 | 117 848 | 117 842 |
| L50 | 2 862 | 1 359 | - | 332 | 236 | 196 | 196 |
| L75 | 8 037 | 4 470 | - | 1 278 | 733 | 587 | 587 |
| Complete BUSCO's (%) | 94.1 | - | - | 98.4 | 98.0 | 98.0 | 98.2 |
| Complete, single copy BUSCO's (%) | 93.5 | - | - | 92.1 | 95.3 | 95.2 | 95.4 |
| Complete, duplicated BUSCO's (%) | 0.6 | - | - | 6.3 | 2.7 | 2.8 | 2.8 |
| Fragmented BUSCO's (%) | 3.9 | - | - | 0.5 | 0.8 | 0.8 | 0.6 |
| Missing BUSCO's (%) | 2.0 | - | - | 1.1 | 1.2 | 1.2 | 1.2 |

Table S7: *Temnothorax americanus* assembly statistics. For further details, see Table S3 legend.

| Assembly | MaSuRCA | HALC-Canu$_{def}$ | Canu$_{def}$-Pilon | Canu$_{err}$-Pilon | PurgeHaplotigs | FinisherSC | Arrow-Pilon |
|---|---|---|---|---|---|---|---|
| No. contigs | 33 648 | 7 960 | 5 585 | 4 202 | 1 829 | 1 229 | 1 229 |
| Total length | 267 639 222 | 370 679 867 | 358 839 550 | 335 049 332 | 287 572 788 | 279 748 504 | 279 543 510 |
| Largest contig | 267 332 | 1 880 024 | 3 579 705 | 4 384 607 | 4 384 607 | 4 384 607 | 4 381 237 |
| GC (%) | 39.08 | 39.19 | 39.14 | 39.21 | 39.21 | 39.20 | 39.20 |
| N50 | 28 826 | 122 674 | 235 295 | 369 006 | 509 480 | 603 326 | 603 026 |
| N75 | 11 469 | 30 921 | 47 521 | 78 171 | 166 114 | 247 685 | 247 480 |
| L50 | 2 514 | 579 | 321 | 199 | 146 | 121 | 121 |
| L75 | 6 147 | 2 256 | 1 158 | 706 | 396 | 298 | 298 |
| Complete BUSCO's (%) | 94.9 | - | 98.1 | 98.2 | 97.7 | 97.8 | 98.2 |
| Complete, single copy BUSCO's (%) | 94.2 | - | 86.0 | 92.0 | 96.1 | 96.2 | 96.9 |
| Complete, duplicated BUSCO's (%) | 0.7 | - | 12.1 | 6.2 | 1.6 | 1.6 | 1.3 |
| Fragmented BUSCO's (%) | 3.6 | - | 0.7 | 0.7 | 0.8 | 0.9 | 0.6 |
| Missing BUSCO's (%) | 1.5 | - | 1.2 | 1.1 | 1.5 | 1.3 | 1.2 |

Table S8: *Temnothorax longispinosus* assembly statistics. For further details, see Table S3 legend.

| Assembly | AllPaths-LG | HALC-Canu$_{def}$ | Canu$_{def}$-Pilon | Canu$_{err}$-Pilon | PurgeHaplotigs$_{err}$ | PurgeHaplotigs$_{def}$ | FinisherSC | Arrow-Pilon |
|---|---|---|---|---|---|---|---|---|
| No contigs | 22 624 | - | 8 013 | 7 938 | 6 402 | 4 012 | 3 430 | 3 430 |
| Total length | 238 236 400 | - | 363 746 299 | 275 125 770 | 257 965 379 | 300 694 620 | 298 389 766 | 298 344 416 |
| Largest contig | 284 717 | - | 1 040 803 | 490 944 | 490 944 | 1 040 803 | 1 040 803 | 1 041 114 |
| GC (%) | 38.78 | - | 38.84 | 39.13 | 39.14 | 38.94 | 38.94 | 38.93 |
| N50 | 30 134 | - | 101 476 | 56 724 | 61 264 | 129 642 | 156 583 | 156 499 |
| N75 | 11 243 | - | 35 819 | 29 170 | 33 104 | 64 750 | 77 995 | 78 015 |
| L50 | 2 153 | - | 918 | 1 412 | 1 267 | 644 | 541 | 541 |
| L75 | 5 361 | - | 2 426 | 3 099 | 2 693 | 1 456 | 1 211 | 1 212 |
| Complete BUSCO's (%) | 95.5 | - | 94.8 | 82.6 | 82.3 | 94.0 | 93.7 | 95.6 |
| Complete, single copy BUSCO's (%) | 94.9 | - | 89.0 | 82.0 | 81.9 | 92.7 | 92.4 | 93.8 |
| Complete, duplicated BUSCO's (%) | 0.6 | - | 5.1 | 0.6 | 0.4 | 1.3 | 1.3 | 1.8 |
| Fragmented BUSCO's (%) | 2.4 | - | 1.8 | 3.5 | 3.6 | 2.1 | 2.0 | 1.1 |
| Missing BUSCO's (%) | 2.1 | - | 3.4 | 13.9 | 14.1 | 3.9 | 4.3 | 3.3 |

Table S9: *Temnothorax rugatulus* assembly statistics. For further details, see Table S3 legend.

| Assembly | MaSuRCA | HALC-Canu$_{def}$ | Canu$_{def}$-Pilon | Canu$_{err}$-Pilon | PurgeHaplotigs | FinisherSC | Arrow-Pilon |
|---|---|---|---|---|---|---|---|
| No contigs | 135 073 | 13 919 | 8 722 | 6 535 | 2 358 | 2 079 | 2 079 |
| Total length | 385 821 052 | 463 678 274 | 467 986 203 | 424 544 191 | 336 328 124 | 335 682 764 | 335 735 536 |
| Largest contig | 352 616 | 731 202 | 3 107 326 | 4 507 738 | 4 507 738 | 4 507 738 | 4 513 070 |
| GC (%) | 39.45 | 38.94 | 39.16 | 39.28 | 39.33 | 39.32 | 39.31 |
| N50 | 7 244 | 53 509 | 135 322 | 235 761 | 425 938 | 450 151 | 450 769 |
| N75 | 2 502 | 25 720 | 35 521 | 46 820 | 118 981 | 140 517 | 141 030 |
| L50 | 12 870 | 1 874 | 584 | 314 | 170 | 155 | 155 |
| L75 | 35 050 | 5 135 | 2 515 | 1 503 | 554 | 494 | 493 |
| Complete BUSCO's (%) | 90.0 | - | 98.1 | 98.5 | 98.4 | 98.3 | 98.4 |
| Complete, single copy BUSCO's (%) | 88.4 | - | 83.9 | 91.8 | 96.0 | 95.7 | 96.1 |
| Complete, duplicated BUSCO's (%) | 1.6 | - | 14.2 | 6.7 | 2.4 | 2.6 | 2.3 |
| Fragmented BUSCO's (%) | 7.3 | - | 0.6 | 0.6 | 0.6 | 0.6 | 0.7 |
| Missing BUSCO's (%) | 2.7 | - | 1.3 | 0.9 | 1.0 | 1.1 | 0.9 |

Table S10: *Temnothorax nylanderi* assembly statistics. For further details, see Table S3 legend.

| Assembly | MaSuRCA | HALC-Canu$_{def}$ | Canu$_{def}$-Pilon | Canu$_{err}$-Pilon | PurgeHaplotigs | FinisherSC | Arrow-Pilon |
|---|---|---|---|---|---|---|---|
| No. contigs | 63 870 | - | 5 108 | 3 357 | 1 785 | 1 555 | 1 555 |
| Total length | 299 554 352 | - | 410 563 087 | 346 999 125 | 312 275 949 | 313 533 164 | 313 389 767 |
| Largest contig | 337 061 | - | 3 912 986 | 4 275 559 | 4 275 559 | 4 919 227 | 4 914 139 |
| GC (%) | 39.24 | - | 38.84 | 38.98 | 39.07 | 39.06 | 39.06 |
| N50 | 21 722 | - | 265 791 | 390 170 | 542 134 | 618 788 | 619 324 |
| N75 | 5 290 | - | 55 617 | 108 502 | 162 861 | 198 066 | 197 671 |
| L50 | 3 035 | - | 289 | 161 | 123 | 111 | 111 |
| L75 | 10 343 | - | 1 194 | 605 | 405 | 349 | 349 |
| Complete BUSCO's (%) | 94.8 | - | 98.4 | 98.3 | 98.3 | 98.2 | 98.5 |
| Complete, single copy BUSCO's (%) | 94.1 | - | 86.8 | 95.1 | 97.1 | 95.7 | 96.1 |
| Complete, duplicated BUSCO's (%) | 0.7 | - | 11.6 | 3.2 | 1.2 | 2.5 | 2.4 |
| Fragmented BUSCO's (%) | 3.1 | - | 0.5 | 0.6 | 0.7 | 0.7 | 0.6 |
| Missing BUSCO's (%) | 2.1 | - | 1.1 | 1.1 | 1.0 | 1.1 | 0.9 |

Table S11: Number of chemosensory receptors in each of the focal species.

| Species | No. *Gr*s | No. *Or*s |
|---|---|---|
| *T. ravouxi* | 49 | 315 |
| *T. unifasciatus* | 117 | 414 |
| *H. sublaevis* | 52 | 309 |
| *L. acervorum* | 127 | 403 |
| *T. americanus* | 41 | 308 |
| *T. longispinosus* | 91 | 419 |
| *T. rugatulus* | 106 | 421 |
| *T. nylanderi* | 128 | 418 |

Table S12: Proportion of chemoreceptor orthologous clusters which exhibited gene losses or gains in each of the focal species. In slave-maker species, the proportion was determined in relation to their respective host and outgroup species, and for hosts in relation to their respective slave-maker and outgroup species.

| Species | Proportion *Or*s lost (%) | Proportion *Or*s gained (%) | Proportion *Gr*s lost (%) | Proportion *Gr*s gained (%) |
|---|---|---|---|---|
| *T. ravouxi* | 27.69 | 6.19 | 47.22 | 1.14 |
| *T. unifasciatus* | 7.49 | 7.82 | 8.33 | 13.89 |
| *H. sublaevis* | 25.08 | 1.30 | 55.56 | 4.17 |
| *L. acervorum* | 3.26 | 7.49 | 0 | 29.17 |
| *T. americanus* | 29.97 | 2.93 | 44.44 | 1.14 |
| *T. longispinosus* | 7.49 | 5.21 | 4.17 | 6.94 |

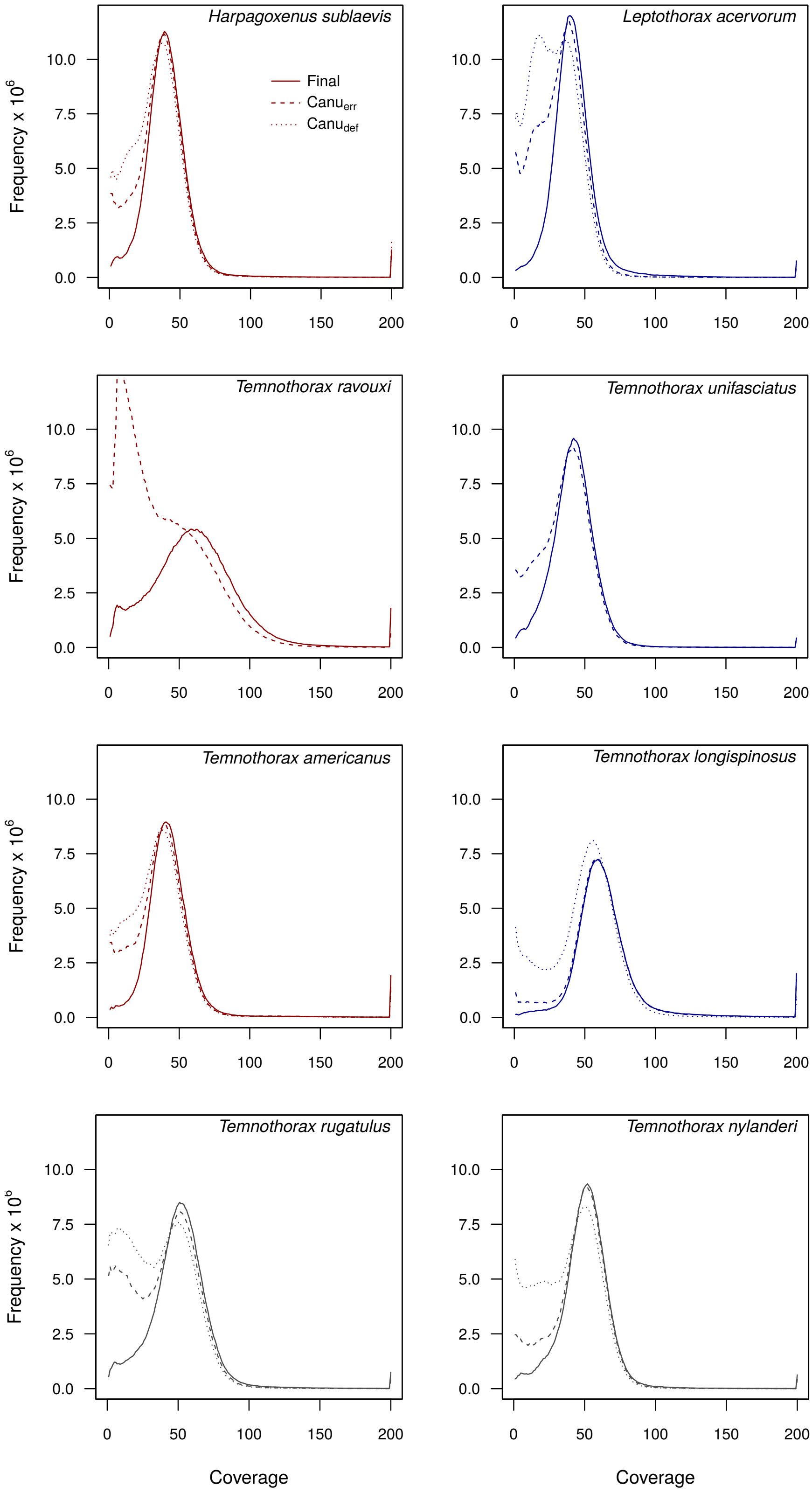


Figure S1: Read depth distribution of the eight genome assemblies.

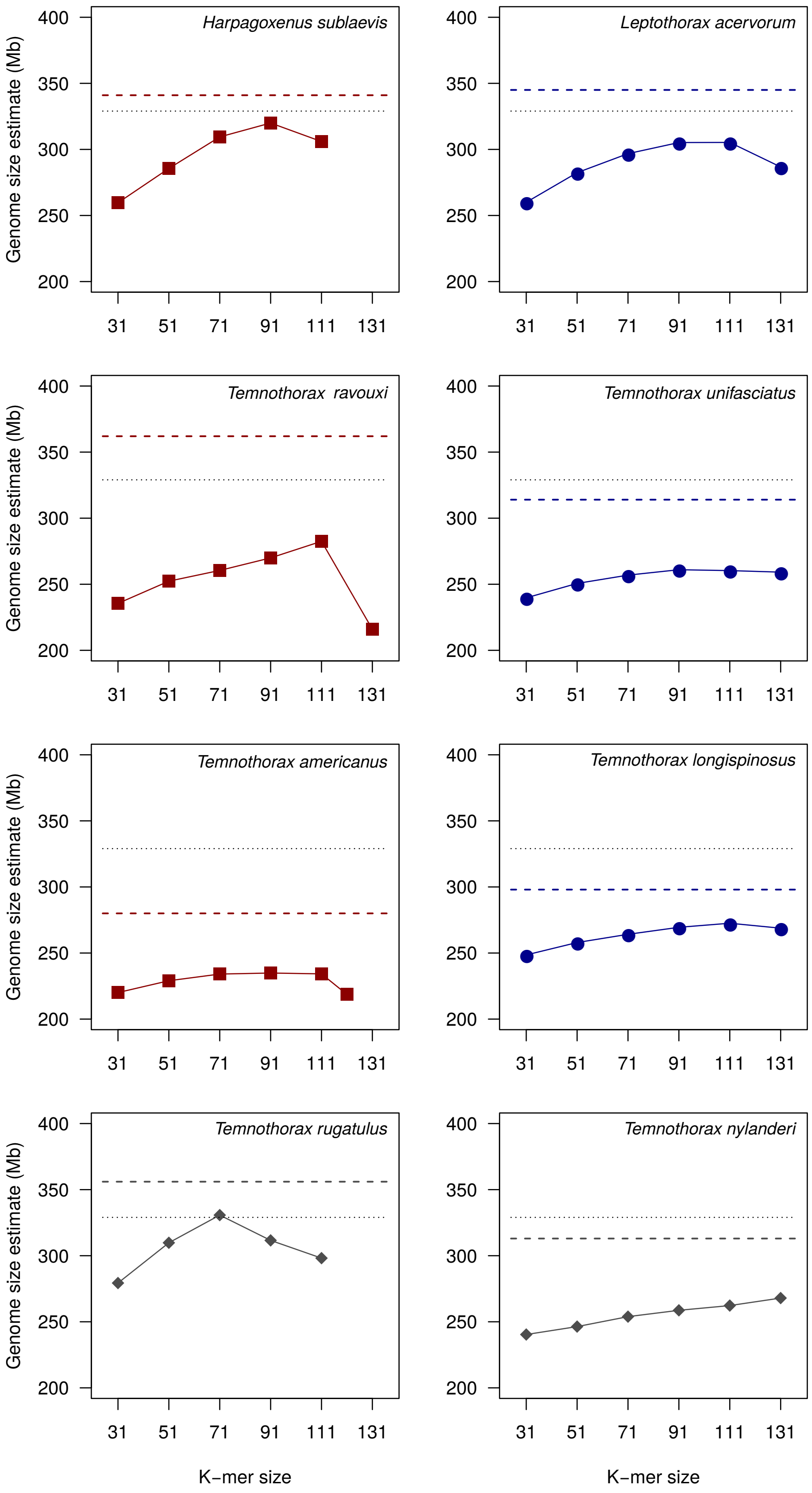


Figure S2: K-mer based genome size estimates. Short dashed, black line represents the average genome size of Myrmicinae [Tsutsui et al., 2008], long-dashed, colored line represents the final, PacBio assembly sizes.

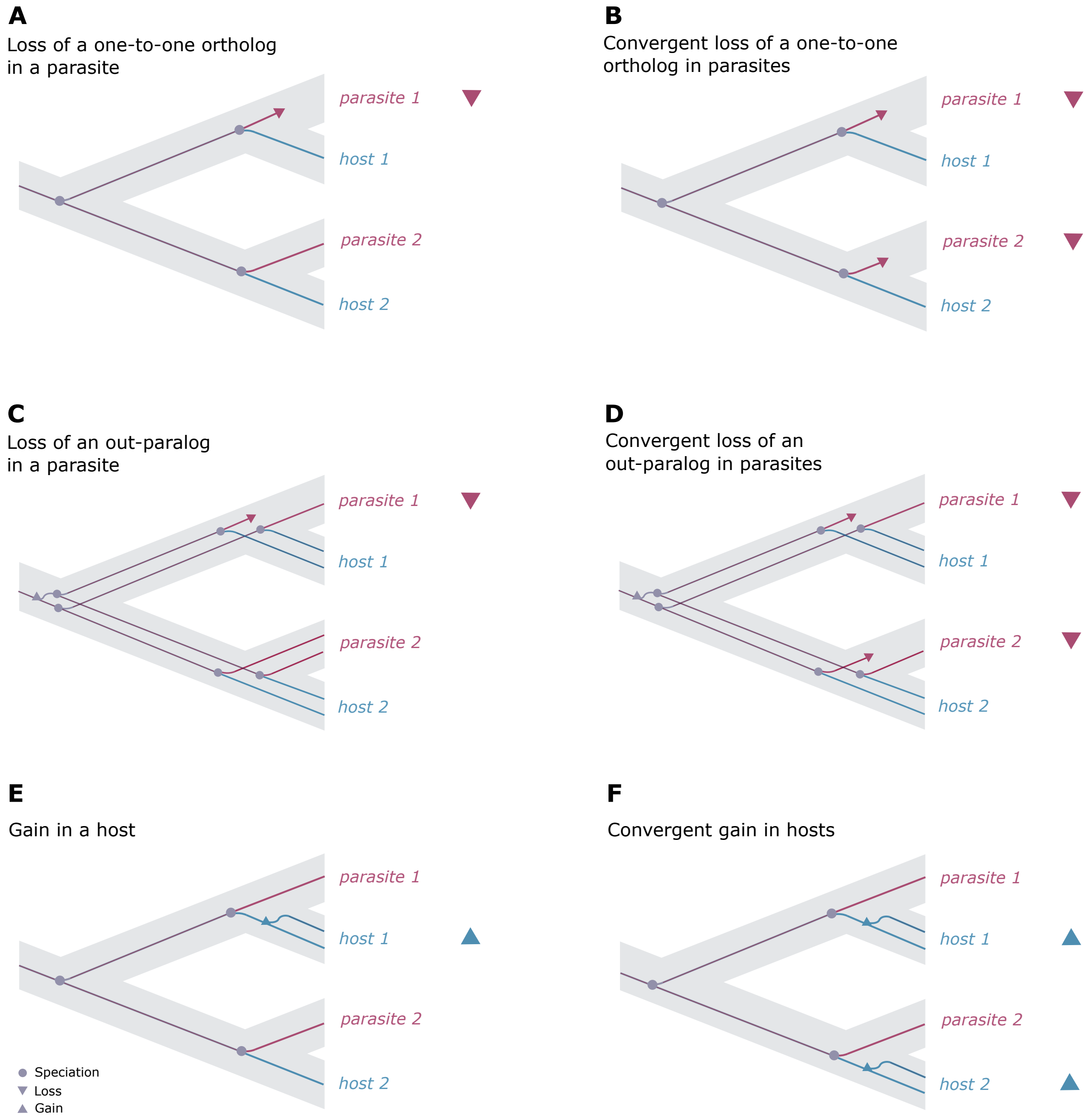


Figure S3: Conceptual diagrams of gene loss and gain in two slave-maker ant (parasite) species and / or hosts. Left panel: loss and gain in individual species, right panel: convergent loss or gain across all parasites or hosts.

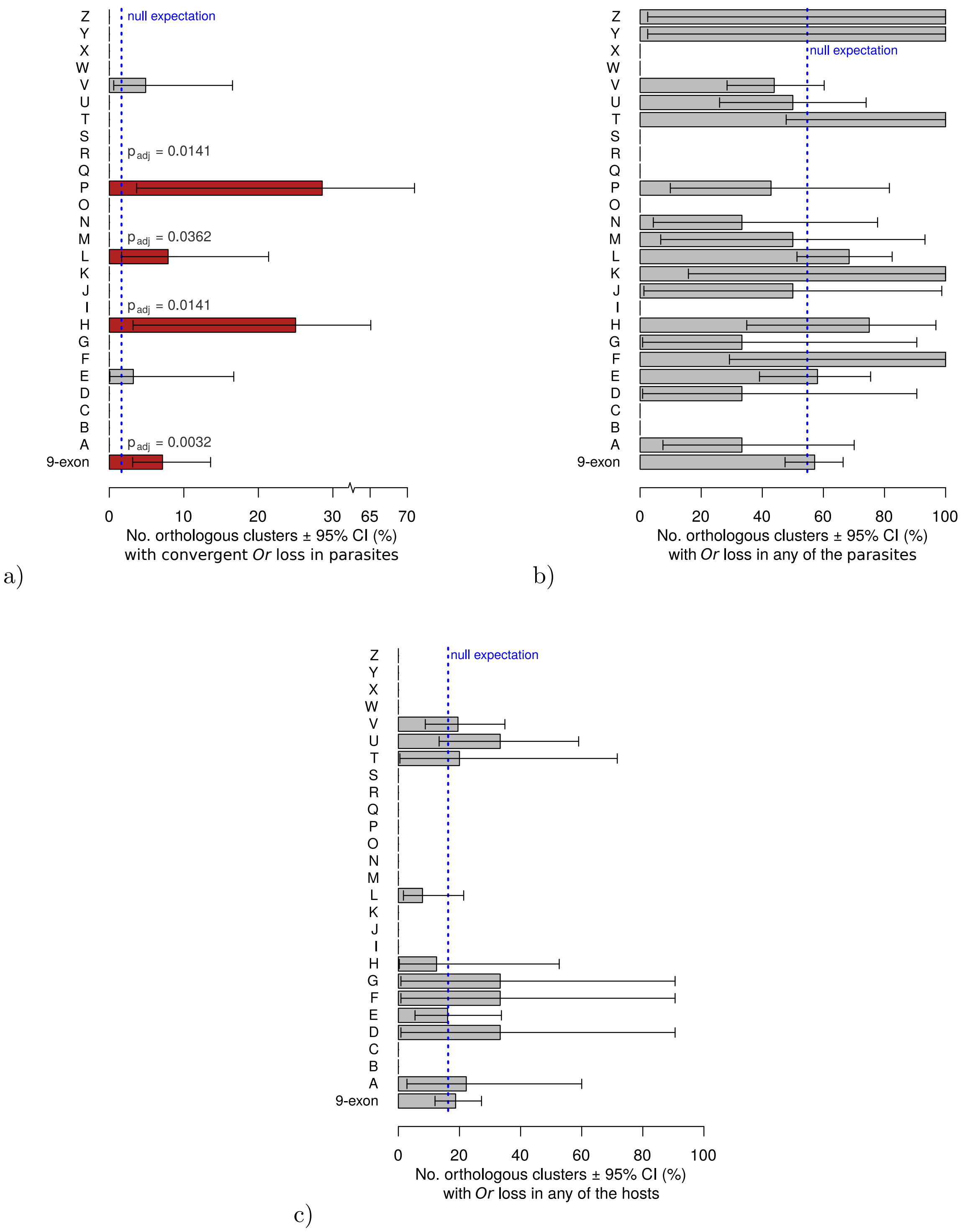


Figure S4: *Or* loss in relation to lifestyle.

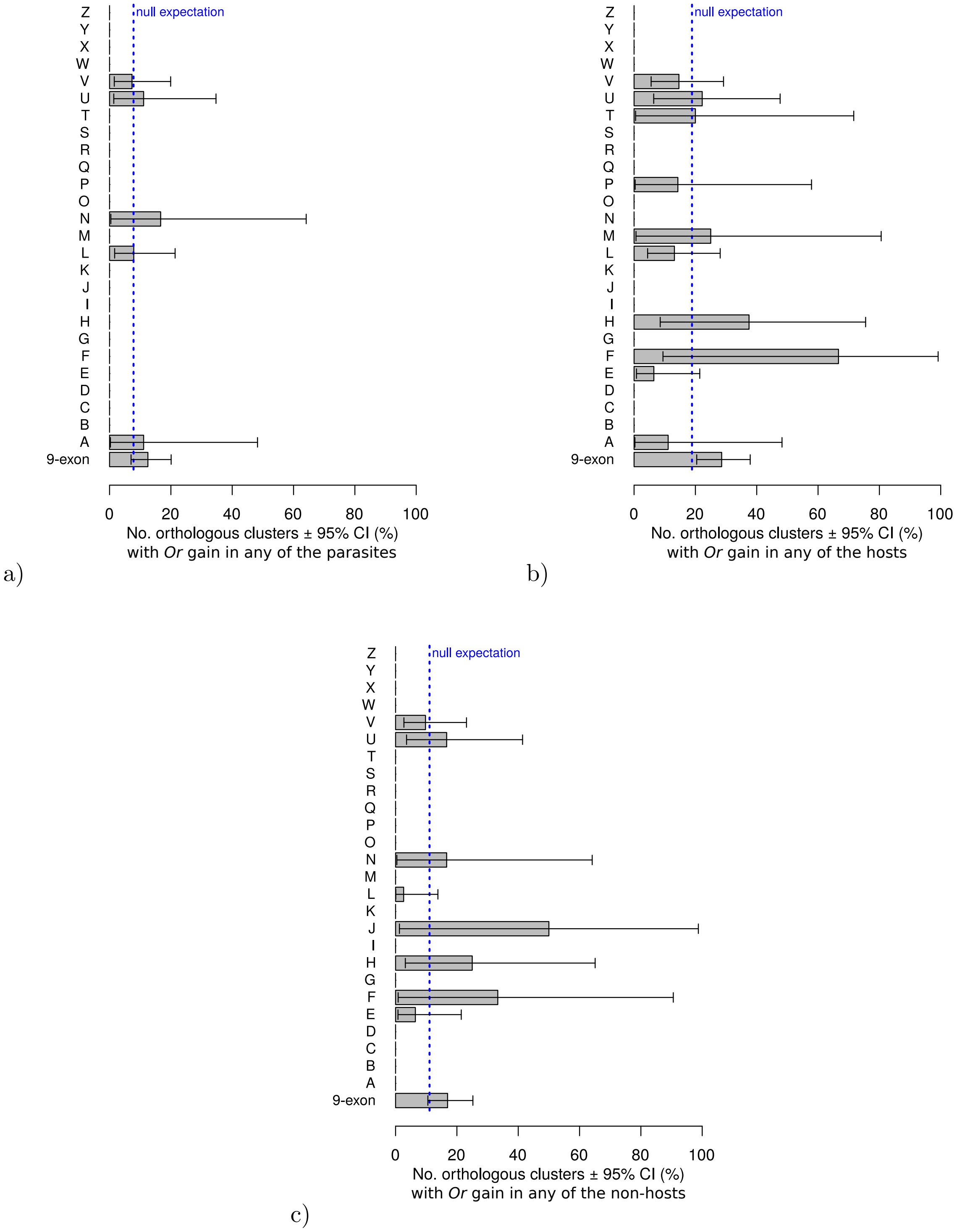


Figure S5: *Or* gain in relation to lifestyle.

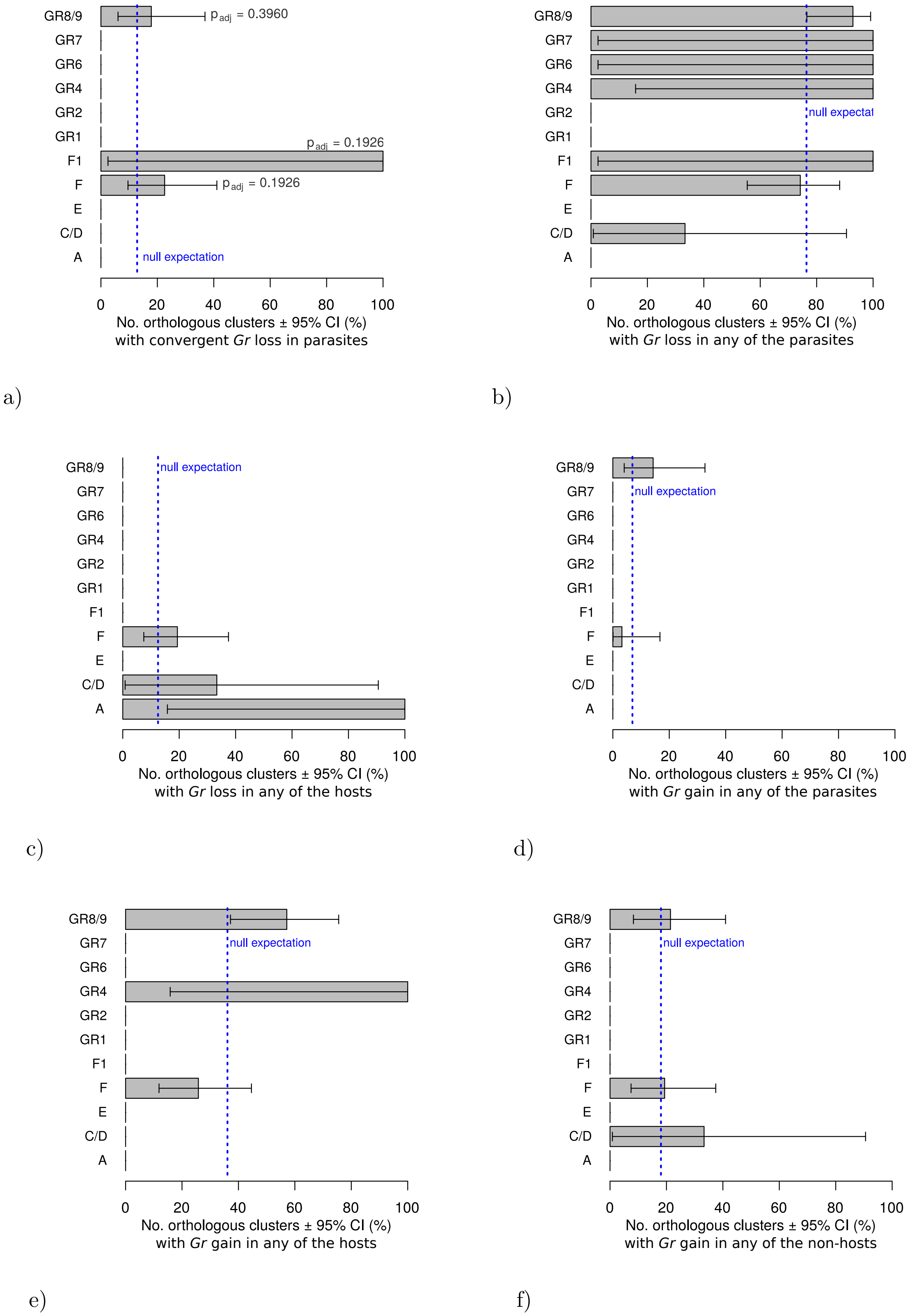


Figure S6: *Gr* gain and loss in relation to lifestyle.

#### References


Ergude Bao and Lingxiao Lan. Halc: High throughput algorithm for long read error correction. BMC Bioinformatics, 18(1):204, Apr 2017. ISSN 1471-2105. doi: 10.1186/s12859-017-1610-3.

Mark J. Chaisson and Glenn Tesler. Mapping single molecule sequencing reads using basic local alignment with successive refinement (blasr): application and theory. BMC Bioinformatics, 13(1):238, Sep 2012. ISSN 1471-2105. doi: 10.1186/1471-2105-13-238.

Alexey Gurevich, Vladislav Saveliev, Nikolay Vyahhi, and Glenn Tesler. Quast: quality assessment tool for genome assemblies. Bioinformatics, 29(8):1072–1075, 2013. doi: 10.1093/bioinformatics/btt086.

Jaime Huerta-Cepas, François Serra, and Peer Bork. ETE 3: Reconstruction, Analysis, and Visualization of Phylogenomic Data. Molecular Biology and Evolution, 33(6):1635–1638, 02 2016. ISSN 0737-4038. doi: 10.1093/molbev/msw046.

Kazutaka Katoh and Daron M. Standley. MAFFT Multiple Sequence Alignment Software Version 7: Improvements in Performance and Usability. Molecular Biology and Evolution, 30(4):772–780, 01 2013. ISSN 0737-4038. doi: 10.1093/molbev/mst010.

S Koren, B. P. Walenz, K. Berlin, J. R. Miller, and A. M. Phillippy. Canu: scalable and accurate long-read assembly via adaptive k-mer weighting and repeat separation. bioRxiv, August 2016.

Ka-Kit Lam, Kurt LaButti, Asif Khalak, and David Tse. Finishersc: a repeat-aware tool for upgrading de novo assembly using long reads. Bioinformatics, 31(19):3207–3209, 2015. doi: 10.1093/bioinformatics/btv280.

Ben Langmead and Steven L. Salzberg. Fast gapped-read alignment with bowtie 2. Nature Methods, 9: 357–359, 2012. doi: 10.1038/nmeth.1923.

Heng Li. Minimap2: pairwise alignment for nucleotide sequences. Bioinformatics, 34(18):3094–3100, 2018. doi: 10.1093/bioinformatics/bty191.

Morgan N. Price, Paramvir S. Dehal, and Adam P. Arkin. Fasttree 2 – approximately maximum-likelihood trees for large alignments. PLOS ONE, 5(3):1–10, 03 2010. doi: 10.1371/journal.pone.0009490.

Michael J Roach, Simon A Schmidt, and Anthony R Borneman. Purge haplotigs: Synteny reduction for third-gen diploid genome assemblies. bioRxiv, 2018. doi: 10.1101/286252.

Leena Salmela and Eric Rivals. Lordec: accurate and efficient long read error correction. Bioinformatics, 30(24):3506–3514, 2014. doi: 10.1093/bioinformatics/btu538.

Felipe A. Simão, Robert M. Waterhouse, Panagiotis Ioannidis, Evgenia V. Kriventseva, and Evgeny M. Zdobnov. Busco: assessing genome assembly and annotation completeness with single-copy orthologs. Bioinformatics, 31(19):3210–3212, 2015. doi: 10.1093/bioinformatics/btv351.

Neil D. Tsutsui, Andrew V. Suarez, Joseph C. Spagna, and J. Spencer Johnston. The evolution of genome size in ants. BMC Evolutionary Biology, 8(1):64, 2008. ISSN 1471-2148. doi: 10.1186/1471-2148-8-64.

Bruce J. Walker, Thomas Abeel, Terrance Shea, Margaret Priest, Amr Abouelliel, Sharadha Sakthikumar, Christina A. Cuomo, Qiandong Zeng, Jennifer Wortman, Sarah K. Young, and Ashlee M. Earl. Pilon: An integrated tool for comprehensive microbial variant detection and genome assembly improvement. PLOS ONE, 9(11):1–14, 11 2014. doi: 10.1371/journal.pone.0112963.

Aleksey V. Zimin, Guillaume Marçais, Daniela Puiu, Michael Roberts, Steven L. Salzberg, and James A. Yorke. The masurca genome assembler. Bioinformatics, 29(21):2669–2677, 2013. doi: 10.1093/bioinformatics/btt476.
